# Asymmetric MukB ATPases are regulated independently by the N- and C-terminal domains of MukF kleisin

**DOI:** 10.1101/180729

**Authors:** Katarzyna Zawadzka, Pawel Zawadzki, Rachel Baker, Karthik V. Rajasekar, David J. Sherratt, Lidia K. Arciszewska

**Author notes:** communicating authors (David J. Sherratt, Lidia K. Arciszewska).

## Abstract

The *Escherichia coli* SMC complex, MukBEF, acts in chromosome segregation. MukBEF shares the distinctive architecture of other SMC complexes, with one prominent difference; unlike other kleisins, MukF forms dimers through its N-terminal domain. We show that a 4-helix bundle adjacent to the MukF dimerization domain interacts functionally with the MukB coiled-coiled ‘neck’ adjacent to the ATPase head, forming an asymmetric tripartite complex, as in other SMC complexes. Since MukF dimerization is preserved during this interaction, MukF directs the formation of dimer of dimers MukBEF complexes, observed previously *in vivo*. The MukF N- and C-terminal domains stimulate ATPase independently and additively, consistent with them each targeting only one of the two MukB ATPase active sites in the asymmetric complex. We demonstrate that MukF interaction with the MukB neck turns over during cycles of ATP binding and hydrolysis *in vivo* and that impairment of this interaction leads to MukBEF release from chromosomes.

## Introduction

SMC (<u>S</u>tructural <u>M</u>aintenance of <u>C</u>hromosomes) complexes have important roles in managing and processing chromosomes in all domains of life (Gligoris and Löwe, 2017; Nolivos and Sherratt, 2015; Uhlmann, 2016). The distinctive architecture of SMC proteins is conserved with the N-and C-terminal globular domains coming together to form an ATPase head and the intervening polypeptide folding upon itself to form ∼50 nm long intramolecular coiled-coil arms, with a dimerization hinge distal from the head (Figure 1). Upon ATP binding, the heads of SMC dimers engage to generate two ATPase active sites (Haering et al., 2002 ***Lemmens et al., 2014***). In eukaryotes, SMC complexes are exclusively heterodimeric, whilst those in bacteria are homodimers. Nevertheless, the distinctive SMC architecture is conserved, with a kleisin protein linking the two ATPase heads of an SMC dimer, thereby forming a large tripartite proteinaceous ring (Figure 1 inset). Essential accessory ‘kite’ or ‘hawk’ proteins bind the kleisin (Palecek and Gruber, 2015; Wells et al., 2017). Hawk proteins are present in cohesins and condensins, while kites are present in bacterial SMC complexes, including MukBEF, and eukaryote SMC5/6 complexes. This suggests that the bacterial complexes are more evolutionarily related to the SMC5/6 complexes of eukaryotes than to eukaryote cohesins and condensins. A substantial body of work has led to the hypothesis that DNA segments are topologically entrapped within these tripartite rings. ATP binding and hydrolysis are required for the entrapping of DNA within the rings, loading, and for DNA release, unloading (Arumugam et al., 2003; Çamdere et al., 2015; Cuylen et al., 2011; Gruber et al., 2003; Haering et al., 2002; Haering et al., 2008; Hu et al., 2011; Kanno et al., 2015; Nasmyth 2001; Nasmyth, 2011; Murayama and Uhlmann, 2014; Uhlmann, 2016; Wilhelm et al., 2015).

**Figure 1.**
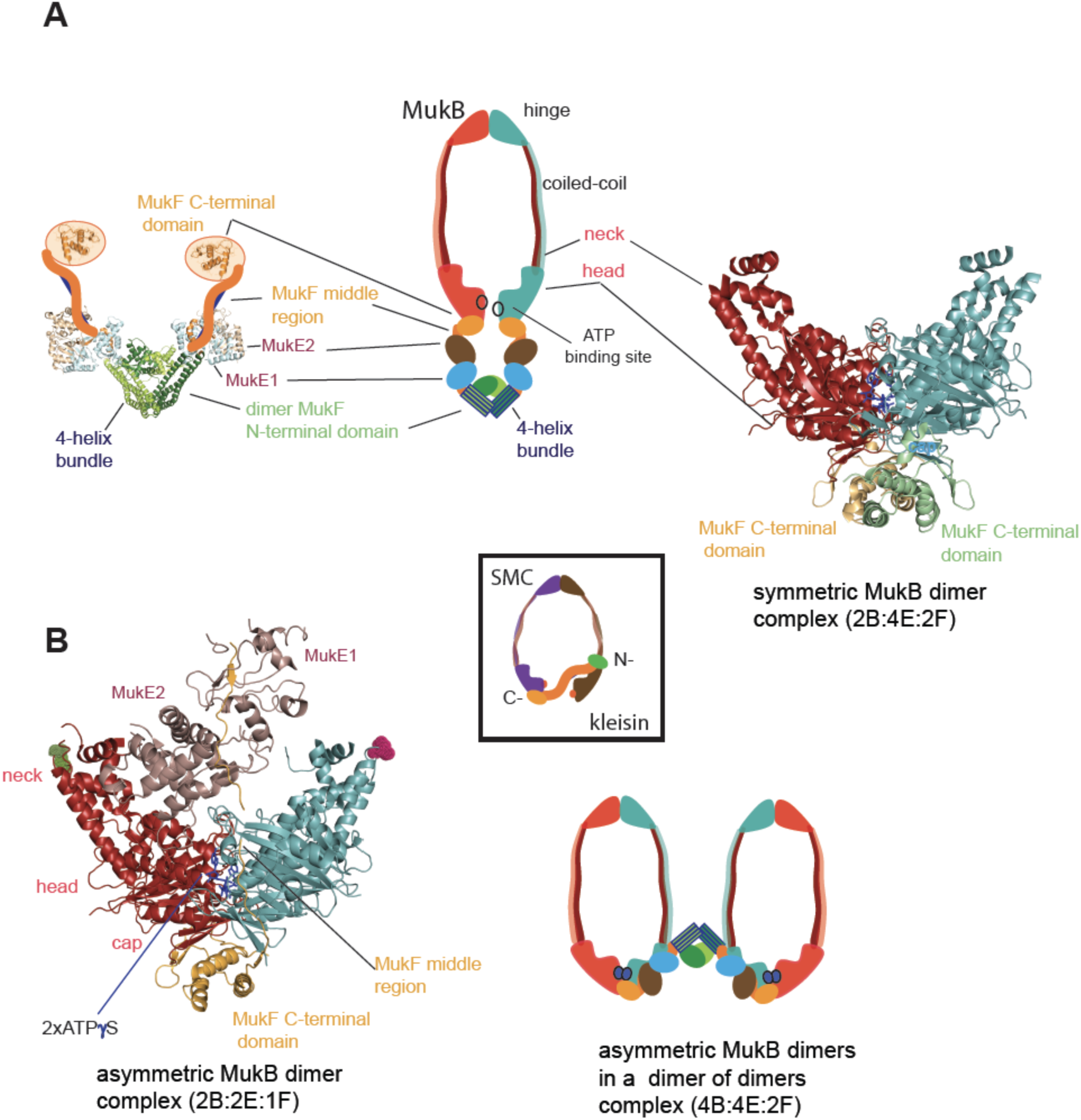
MukBEF complexes. (A) middle panel: classical view of the proposed symmetric complex of MukBEF in the absence of ATP, with a molecular ratio of 2B:4E:2F. Two monomers of a MukB dimer are joined by their hinge domains. A dimer of MukF, decorated with four MukE monomers, interacts through its C-terminal domains with the cap regions of the MukB heads. Left panel: schematic of a MukF dimer based on the structure of the MukF N-terminal region bound by MukE (pdb, 3EUH; Woo et al., 2009) and the C-terminal domain structure (pdb 3EUK, Woo et al., 2009), with a cartoon of the intervening middle region. Right panel: crystal structure of symmetric complex of *H.ducreyi* hMukE-hMukF(M+C)-hMukBhd^EQ^-ATP-γS obtained in the presence of DNA (pdb 3EUH, Woo et al., 2009); the symmetric complex shows two heads (red and teal) bound by two MukF C-terminal domains (yellow and green); the heads are close together but not in engaged conformation. Two molecules of ATP-γS are bound at their interface. MukE and MukF middle regions were not observed in the structure. (B) left: crystal structure of *H.ducreyi* hMukE-hMukF(M+C)-hMukBhd^EQ^-ATP-γS asymmetric complex (pdb 3EUK, Woo et al., 2009) obtained in the absence of DNA. The asymmetric complex is formed by ATP binding-mediated head engagement; one MukFE2 is expelled from the MukB head complex and the molecular ratio is 2B:2E:1F; the residues adjacent to the neck coiled coil regions, which are not in the structure (at the coiled-coil exit points), are indicated on each head by green and pink dots, respectively. The C-terminal domain of MukF, yellow, binds to a cap region of one monomer of MukB, while the C-terminus proximal part of the MukF middle region interacts with the head of the partner MukB monomer. Muk E dimer bound to a remaining, N-terminus proximal part of the MukF linker is shown in brown; right: a cartoon of MukBEF dimer of dimers with stoichiometry of 4B:4E:2F, which was inferred from *in vivo* stoichiometry measurements (Badrinarayanan et al., 2012a). Inset centre shows a cartoon of a typical SMC-kleisin tripartite ring.

*E. coli* and its closest γ-proteobacterial relatives, encode an apparently distant SMC relative, MukBEF, with little primary sequence homology to other SMCs (***Nolivos and Sherratt, 2014***). Organisms encoding MukBEF have co-evolved a number of other distinctive proteins, some of which interact with MukB physically and/or functionally; specifically, topoisomerase IV and MatP both interact with MukB *in vitro* and *in vivo* (Brézellec et al., 2006; Hayama and Marians, 2010; Hayama et al., 2013; Li et al., 2010; Nicolas et al., 2016; Nolivos et al., 2016; Vos et al., 2013). MukB forms SMC homodimers, whereas MukF is the kleisin and MukE the kite protein that binds MukF (Palecek and Gruber, 2015). All three proteins of the MukBEF complex are required for function and their impairment leads to defects in chromosome segregation, manifested by impairment of segregation of newly replicated origins (*ori*) and mis-orientation of chromosomes with respect to their genetic map within cells (Danilova et al., 2007). In rich media, this leads to inviability at higher temperatures and formation of anucleate cells during permissive low-temperature growth (Niki et al., 1991; Yamanaka et al., 1996).

Where characterized, most SMC complexes bind their cognate monomeric kleisins asymmetrically, with their N-terminal regions binding the SMC ‘neck’ adjacent to the ATPase head of the molecule distal to the molecule binding the kleisin C-terminus, thereby-forming the tripartite protein ring (Figure 1 centre inset) (Bürmann et al., 2013; Gligoris et al., 2014; Gligoris and Löwe, 2017; Gruber et al., 2003; Haering et al., 2004; Huis in’t Veld et al., 2014). MukF is an atypical kleisin, in that it forms stable dimers through interacting N-terminal winged-helix domains (WHD), while its C-terminal domain interacts with the MukB head at the cap, as is the case for characterized SMC kleisins (Fennell-Fezzie et al., 2005, Woo et al., 2009). Therefore, one MukB dimer is expected to bind a MukF dimer and two MukE dimers in the absence of ATP (Figure 1A centre); ATP binding leads to head dimerization (Figure 1A right panel), accompanied by steric expulsion of one of the two MukF C-terminal domains and head engagement (Woo et al., 2009, Figure 1B left panel).

Here, we reveal that MukF, like other characterized kleisins, interacts functionally with the SMC MukB neck, through a 4-helix bundle in its N-terminal domain, while its C-terminal domain interacts with the MukB head at the cap. We show that this interaction with the MukB neck is required for MukBEF function *in vivo,* and infer that this interaction is established and broken during cycles of ATP binding and hydrolysis. Impairment of this interaction *in vivo* leads to release of MukBEF clusters from chromosomes. Interactions of the MukF N-terminal domain with the MukB neck and MukF C-terminal domain with the MukB head, activate MukB ATPase independently and additively *in vitro*, with the addition of both fragments restoring wild type MukB ATPase levels. Each of these ATPase activities was inhibited by MukE, with the inhibition being relieved in the presence of DNA. We show that interaction of the MukF N-terminal domain with the MukB neck did not compromise MukF dimerization. Therefore, MukF dimerization in heads-engaged MukBEF complexes directs formation of dimers of MukBEF dimers, thereby explaining the stoichiometry observed *in vivo* (Badrinarayanan et al., 2012, Figure 1B right panel).

## Results

### The MukF N-terminal domain interacts with the MukB neck

Because of the intriguing distinction between dimeric MukF and monomeric kleisins (Figure 1), we set out to test, if the MukF N-terminal domain would also interact with the MukB neck, thereby exhibiting the architecture of other SMC dimers and their cognate kleisins. In order to undertake an initial characterization of MukB-MukF interactions, C- and N-terminal MukF Flag-tagged truncations, immobilized on anti-Flag resin, were analyzed for binding to MukB derivatives. Intact MukB and truncated derivatives containing just the ATPase head (MukB_H_), or the head plus a segment of adjacent ‘neck’ coiled-coil (MukB_HN_), which would be expected to contain any interaction determinants for MukF, were used (Figure 2A; Bürmann et al., 2013, Gligoris et al., 2014). MukF C-terminal derivatives, FC1 and FC2, interacted with MukB, and all of its derivatives, as expected, because the MukB ATPase head participates in this interaction (Figure 2B; Figure 1; Woo et al., 2009). FN2, containing the N-terminal dimerization WHD and an adjacent 4-helix bundle, interacted strongly with intact MukB and MukB_HN_, but not with MukB_H_, consistent with FN2 interacting with the MukB neck, but not the head (Figure 2B). FN1, containing just the WHD involved in dimerization showed no interactions with the MukB derivatives, while FN3, containing the 4-helix bundle (helices 6-9), and FN4, carrying only helices 8 and 9, interacted with MukB and MukB_HN_, but not MukB_H_. Consistent with this, FN6 lacking helices 8 and 9 failed to show an interaction in SEC-MALS assays (below) and FN7, lacking helix 9 failed to interact with MukB_HN_ (Figure 2B bottom right panel). We conclude that helices 8 and 9 of the 4-helix bundle are sufficient for interaction with the MukB neck. Helix 9 is essential either because it interacts with the neck, or because it is required for proper folding of the helix 8-9 polypeptide.

**Figure 2.**
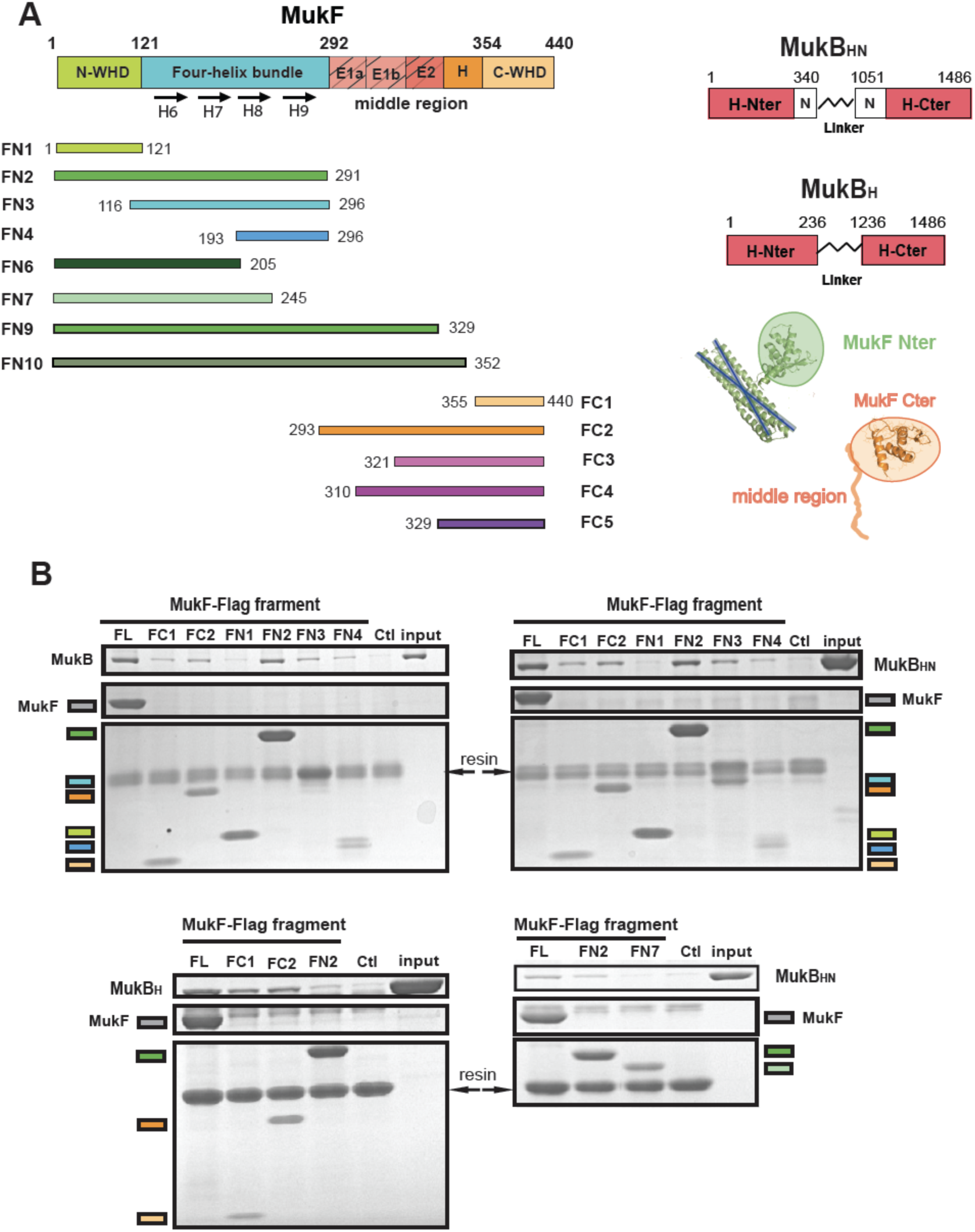
The MukF N-terminal domain interacts with MukB neck. (A, left panel) Schematics of the MukF truncations. The MukF N-terminal WHD is responsible for MukF dimerization, while the C-terminal WHD interacts with the MukB head (Fennell-Fezzie et al., 2005, Woo et al., 2009) The middle region contains binding sites for the MukE dimer (E1, E2,) and the C-terminal part of the extended polypeptide that interacts with the MukB engaged head (‘H’; Woo et al., 2009). Monomer 1 of the MukE dimer binds helical E1a and part of the acidic linker E1b, while the second MukE monomer, within the dimer binds E2, also part of the linker. (A, right panel) The MukB head variant (MukB_H_) carries N- and C-terminal regions that together constitute head domain, joined by 18 aa residue flexible linker, while MukB ‘head and neck’ variant (MukB_HN_) in addition carries the predicted, head proximal (C0) segment of coiled-coil (Weitzel et al., 2011) 104/185 amino acid residues adjacent to MukB N- and C-terminal domains, respectively. Cartoons of structures of MukF N- and C-terminal domains are included. (B) Pull-down assay using MukF-FLAG tagged fragments as baits for the indicated MukB derivatives. The amounts of recovered MukB, MukB_HN_ or MukB_H_ are shown within the top boxed portion of the gel in each panel, alongside the MukB derivative input and Ctl., a control with no added bait.

To confirm these observations, and to determine the molecular mass of the complexes, we used size exclusion chromatography-multi-angle light scattering (SEC-MALS; Figure 3). MukB_HN_ was a monomer in solution, while FN2 was dimeric, as expected from structural analyses (Fennell-Fezzie et al., 2005, Woo et al., 2009). When mixed at a molar ratio of 1 MukB_HN_ monomer:1.25 FN2 dimers, two additional peaks of masses 165kDa and 284kDa were evident in addition to the MukB_HN_ monomers (Figure 3A left panel). We interpret these as complexes in which either one or two MukB_HN_ molecules bound independently to a single FN2 dimer. Consistent with this interpretation, more of the larger complexes were observed at higher MukB_HN_ to 2FN2 ratios (3:1; Figure 3-figure supplement 1). Therefore, the interaction between the MukF N-terminal domain and the MukB neck does not compromise MukF dimerization. The relatively low proportion of complexes of stoichiometry MukB_HN_-2FN2-MukB_HN_ as compared to MukB_HN_-2FN2 in the presence of a large excess of MukB_HN_, indicates that binding of the second MukB_HN_ to MukB_HN_-2FN2 complex may be less favourable than binding of the first MukB_HN_ to 2FN2.

**Figure 3.**
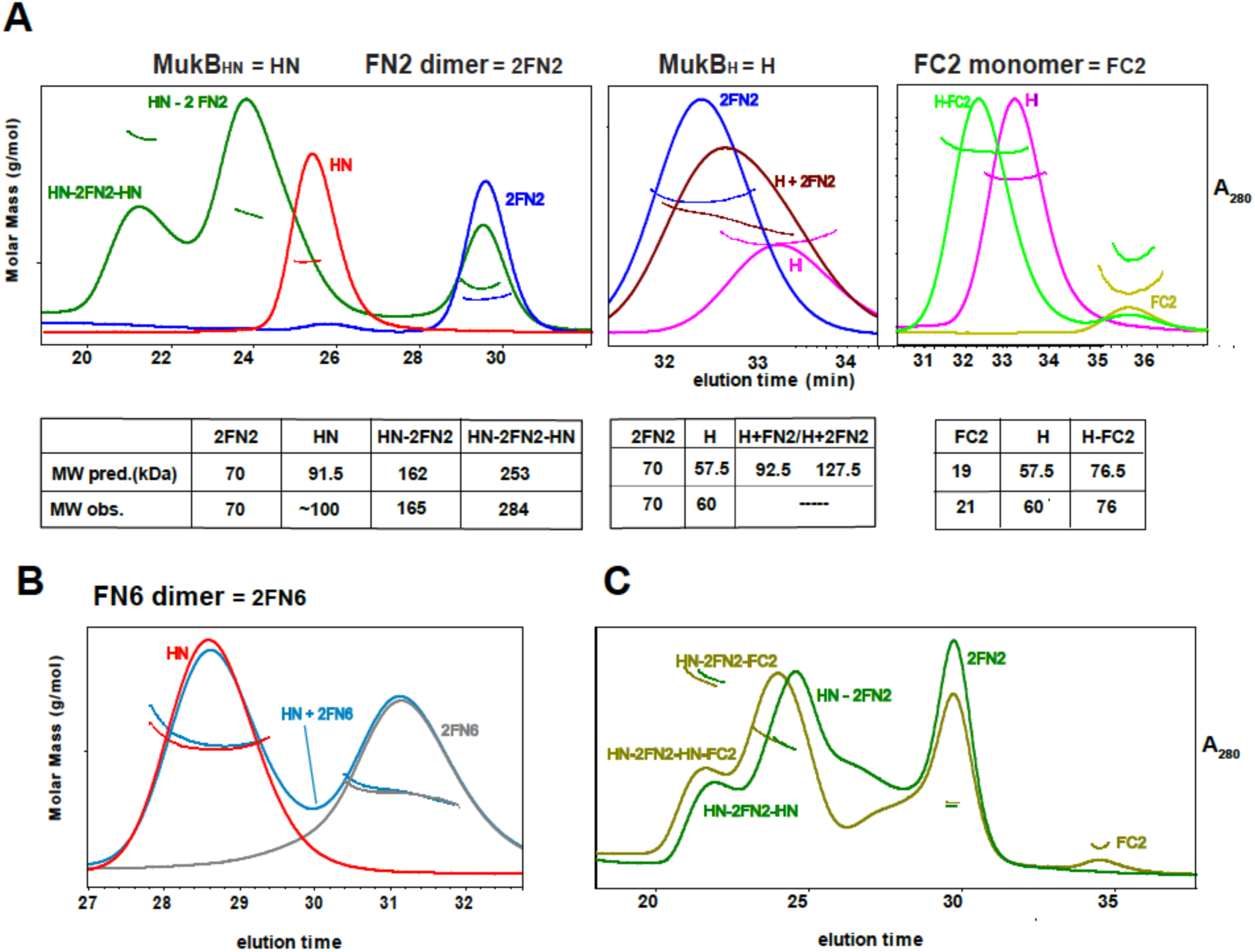
Complexes of MukF N- and C-terminal domains with MukB head variants. Binding and stoichiometry of complexes was determined by SEC-MALS analyses. (A) left panel: MukB_HN_ (red), 2FN2 (blue), and MukB_HN_+2FN2 (green) at 1:1.25 monomer:dimer (1:2.5 m:m) molar ratio; middle panel: MukB_H_ (pink), 2FN2 (blue), and MukB_H_+2FN2 (brown) at 1:0,25 m:d (1:0.5 m:m) ratio; right panel: MukB_H_ (pink), FC2 (lime green), and MukB_H_+FC2 (green) at 1:1 m:m ratio (B) MukB_HN_ (red), 2FN6 (grey), and MukB_HN_+2FN6 (blue) at 1:0.25 m:d (1:0.5 m:m) ratio. (C) MukB_HN_+2FN2 at 1:1 m:d (1:2 m:m) ratio (dark green), and MukB_HN_+2FN2+FC2 at 1:1:1 m:d:m (1:2:1 m:m:m) ratio (olive green).

In agreement with the Flag-MukF-MukB interaction assays, FC2, but not FN2, formed complexes with MukB_H_ (Figure 3A middle and right panels). FN6, which lacks the two C-terminal helices, 8 and 9, of the 4-helix bundle, failed to form complexes with MukB_HN_ (Figure 3B). Addition of ATP did not significantly alter the nature or abundance of complexes containing MukB_HN_ and FN2 or FC2 (Figure 3-figure supplement 2). This is consistent with MukB_HN_, which is a monomer in solution, being unable to form stable heads-engaged dimers with either FN2 or FC2 in the presence of ATP.

We next tested whether monomers of MukB_HN_ can simultaneously bind both FN2 and FC2. SEC analysis (Figure 3C) showed that mixtures of MukB_HN_, FN2 and FC2 yielded larger complexes (olive green trace) than those formed with MukB_HN_ and FN2 alone (dark green trace), consistent with binding of both FN2 and FC2 to a single monomer of MukB_HN_. Nevertheless, it was not possible to assign precise masses to these by light scattering, because of the dynamic nature of the complexes and an inability to completely resolve them under a range of SEC conditions. Therefore, a complex containing a MukB dimer with unengaged heads, bound to a MukF dimer may be stabilized by MukF interactions to both the MukB heads at caps and necks. An equivalent result was observed with *B. subtilis* SMC complexes, with both head and neck of a single SMC molecule being bound simultaneously by kleisin N-and C-terminal domains (Bürmann et al., 2013).

To characterize further the interaction of MukF N- and C-terminal domains to MukB, we determined the binding affinities of fluorescently labelled FN2, FN10, FN3 and FC2 using Fluorescence Correlation Spectroscopy (FCS) and Fluorescence Polarization Anisotropy (FPA). Both domains bound to MukB with similar affinities, with Kds in the 9-26 nM range, suggesting that interactions of the N-terminal and C-terminal MukF domains with the MukB neck and head, respectively, are similarly strong (Figure 3-figure supplement 3). FN10, which in addition to the N-terminal domain also carries the MukF middle region, bound more tightly to MukB consistent with the MukF middle region interacting directly with MukB (see below).

### The MukF C- and N-terminal domains activate MukB ATPase independently and additively

MukB dimers alone had negligible ATPase activity (Figure 4), in agreement with previous reports (Woo et al., 2009). Addition of MukF kleisin led to robust MukB ATPase. The steady state ATPase rate was ∼21 ATP molecules hydrolysed/min/MukB dimer, under conditions of MukF excess (Figure 4-figure supplement 1A). MukF alone did not exhibit ATPase activity. To dissect the MukF requirements for MukB ATPase, we assayed MukF truncations containing either the N-terminal domain (FN2) or the C-terminal domain (FC2) (Figure 2A). Both variants stimulated MukB ATPase independently (Figure 4). Saturating FC2, at a 2.5-fold molar excess, gave 60% of the maximal ATPase obtained with MukF, while saturating FN2 (at a 2.5-fold molar excess) gave 33% of maximum ATPase (Figure 4-figure supplement 1BC). Addition of FN2 and FC2 together restored ATPase to the level observed with wild type MukF. FN6, lacking helices 8 and 9 did not stimulate MukB ATPase, consistent with its failure to interact with the MukB neck (Figure 4-figure supplement 1D).

**Figure 4.**
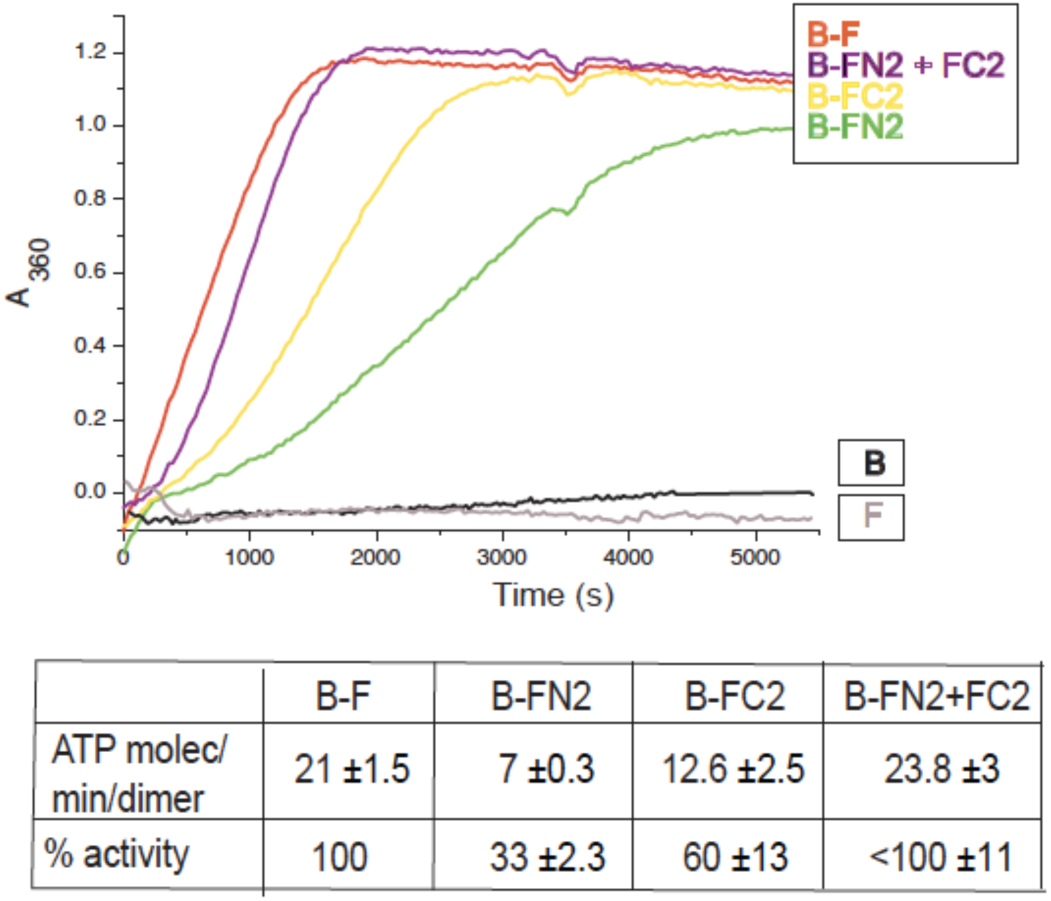
MukF N- and C-terminal domains stimulate MukB ATPase. ATPase activity was measured at concentrations of MukB, 0.5μM MukF/FN2/FC2 1.25μM, i.e. at molar ratio of B:F, 0.5:1.25, m:m. The curves in the graph represent a single experiment; averages of initial rates +/- SD from three experiments are tabulated beneath.

Taken together, the results show that MukB ATPase is activated additively and independently by the N-and C-terminal domains of MukF. Since these domains interact with different monomers in an engaged head MukB dimer, with the MukF C-terminal domain interacting with the head of one MukB monomer, and the N-terminal MukF domain interacting with the neck of the other monomer (Figure1; Woo et al., 2009); we propose that this asymmetric complex contains two independent ATPase domains, one activated by the MukF N-terminal domain and the other activated by the C-terminal domain.

### Characterization of the interactions between the MukB neck and MukF

To gain further insight into the interaction of the MukF 4-helix bundle and the MukB neck, variants altered in the MukB neck and MukF helix 9 were analyzed for their activity and binding. The mutagenesis strategy was informed by MukB and MukF sequence homology and structures of comparable kleisin and SMC neck interactions with kleisin in cohesin, and with *B. subtilis* SMC complexes (Gligoris et al., 2014; Huis in’t Veld et al, 2014, Bürmann et al., 2013). Three triple substitutions in helix 9 of FN2 exhibited an impaired ability to activate MukB ATPase. FN2m2 (with substitutions R279E K283A R286A, red trace) displayed a ∼10-fold reduction in the ability to activate MukB ATPase, while FN2m1 (D272K I275K R279D, yellow trace) and FN2m3 (D261K S265K Q268A, purple trace), showed a ∼2-fold reduction (Figure 5A), Consistent with this, SEC analysis showed that FN2m1 and FN2m2 failed to interact with MukB_HN_ (Figure 5-figure supplement 1); thereby confirming the importance of helix 9 in the interaction with the MukB neck.

**Figure 5.**
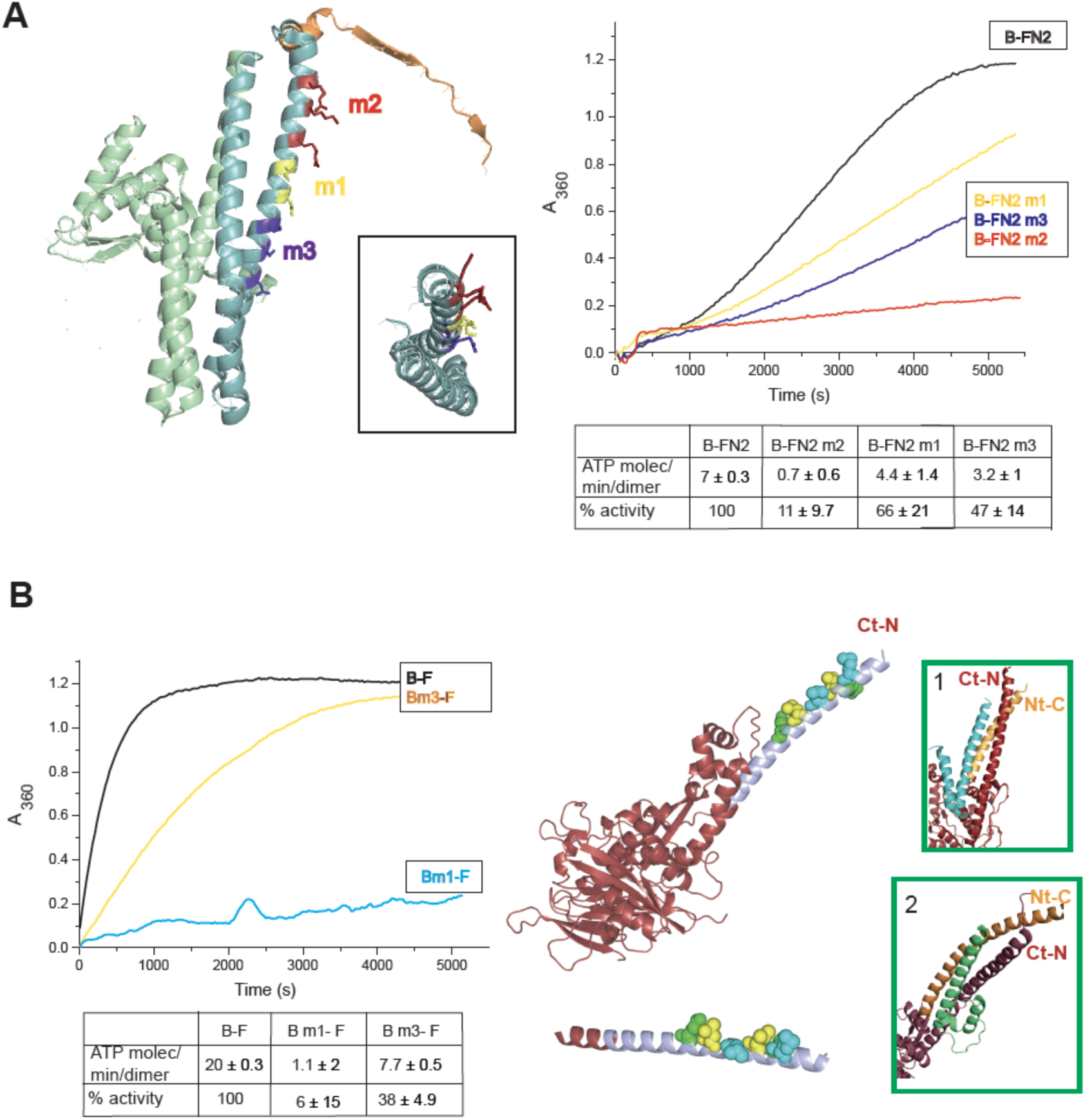
Interface between MukB neck and MukF four helix bundle. (A) left panel: a cartoon of MukF N-terminal domain fragment carrying N-terminal domain (green) and part of the middle region (orange; note that the middle region is not present in FN2). Helices 8 and 9 are indicated in teal, the mutated amino acid residues in mutant variants FN2m1, FN2m2 and FN2m3 are indicated in yellow, red and purple, respectively (residue R279 was altered in both m1 and m2 – here it is included in m2); inset: a view of the two helices from the top. right panel: ATPase activity of the mutated variants; averages of initial rates measurements from 3 experiments are tabulated underneath. (B) ATPase activity of MukB and MukB variants mutated at the neck, MukBm1 blue, and MukBm3 yellow, in the presence of MukF. Averages of initial rates measurements from three experiments are tabulated underneath. MukBm2 had ATPase activity of wt MukB (not shown). A Pymol structure model of MukB head neck variant based on published structure of hMukE-hMukF(M+C)-(hMukBhd^EQ^-ATP-γS). The most N-terminal part of the helix that belongs to the C-terminal subdomain part of the head, and constitutes the neck has been modeled (coloured in lilac) in order to show positions of the mutated residues, which are indicated as follows: Bm1 blue, Bm2 green, and Bm3 orange; below, enlarged view of the neck shown. Insets: Pymol cartoons showing published interactions between kleisin N-terminal domains with SMC necks. 1. *B. subtilis* SMC-ScpA^N^; Bürmann et al., (2013), 2. Smc3-Scc1N; Gligoris et al., (2014). In the neck regions, C-terminal subdomain helices are indicated in dark red, N-terminal domain helices in orange, kleisin helices are indicated in cyan-ScpA and green-Scc1. Ct-N defines the most N-terminal residue of the C-terminal subdomain of the head and Nt-C defines the most C-terminal domain of the N-terminal subdomain in a given SMC head variant.

In addition, we analyzed MukB variants carrying three double amino acid substitutions in the neck, located near MukB C-terminal head domain. They were designed to be on different faces of a putative candidate coiled-coil helix. MukBm1 (M1215K L1222K, blue trace) had <10% of MukB ATPase, MukBm3 (L1219K L1226K, yellow trace) had about 35 % activity while MukBm2 (E1216A E1230A, showed no reduction in ATPase (Figure 5B). Furthermore, MukBm1 and MukBm3 failed to bind FN2 in and FPA assays (Figure 5-figure supplement 2). Consistent with these results, expression *in vivo* of MukBm1 failed to complement the temperature-sensitive growth defect of *mukB* deletion cells, while MukBm2 expression complemented the Muk^-^ phenotype (Figure 5-figure supplement 3). Expression of MukBm3 partially complemented the Muk^-^ phenotype.

Using the *E.coli* MukEF crystal structure (pdb, 3EUH; Woo et al., 2009) along with the structure of the engaged MukB heads from *H.ducreyi* (pdb 3EUK, Woo et al., 2009), we modelled the structure of FN2 dimer bound by two monomers of MukB_HN_. This indicated that unless a major conformational change within FN2 dimer takes place upon MukB_HN_ binding, the arrangement of the heads, imposed by interaction of their necks with 4-helix bundles of the dimer would lead to a different arrangement of the heads to the one revealed by the structure of the engaged MukB heads complex (Woo et al., 2009; Figure 5-figure supplement 4). The motifs that compose the two ATPase active sites in each head monomer are rotated away from each other and therefore distant. Therefore, if simultaneous binding of the two necks within the intact MukB dimer by the N-terminal domains of MukF dimer is possible, it would produce a complex with ‘inactive’ heads arrangement. Whether such complex is generated at any stage of MukBEF activity cycle remains to be determined.

### MukE inhibits MukBF ATPase

MukE showed a concentration-dependent inhibition of MukF-activated MukB ATPase (Figure 6A; Figure 6-figure supplement 1), in agreement with a report by Bahng et al. (2016), which cited our previously unpublished observation. We then tested if MukE could equally inhibit the ATPase activated by the isolated C-and N-terminal domains of MukF. The binding of MukE to a MukBF complex depends on the asymmetric binding of a MukE dimer to the MukF middle region, which also interacts with MukB head in the engaged MukB heads complex (Figure 1B left panel). We therefore, analyzed activity of a series of MukF N-terminal and C-terminal variants that carried different truncations in the middle region (Figure 7A). In the absence of MukE, the N-terminal and C-terminal variants stimulated MukB ATPase at levels that related to the extent of the middle region they contained, presumably due to differences in the stability of their complexes with MukB (Figure 7B). Significantly, FN10 and FC2, both of which carry the entire middle region were equally effective in activating MukB ATPase, thus the MukF N-and C-terminal domains have comparable abilities to activate MukB ATPase. MukE inhibited MukB ATPase activated by the MukF N- and C-terminal domain variants that carried complete MukE dimer binding sites (FN9, FN10 and FC2), with a ∼4-fold greater inhibition of activation by FN10, as compared to FC2 (Figure 7C). MukE was unable to inhibit ATPase stimulated by FN2 and FC5, both of which were lacking MukE binding sites and FC3, which carried only the single E2 MukE binding site. The effect of MukE on FC4, lacking the N-terminal part of the MukE1 binding site (E1A) was to partially inhibit ATPase activation. These data demonstrate that each of the MukB ATPase activities, stimulated independently by the N- and C-terminal domains of MukF can be inhibited by MukE binding to MukF.

**Figure 6.**
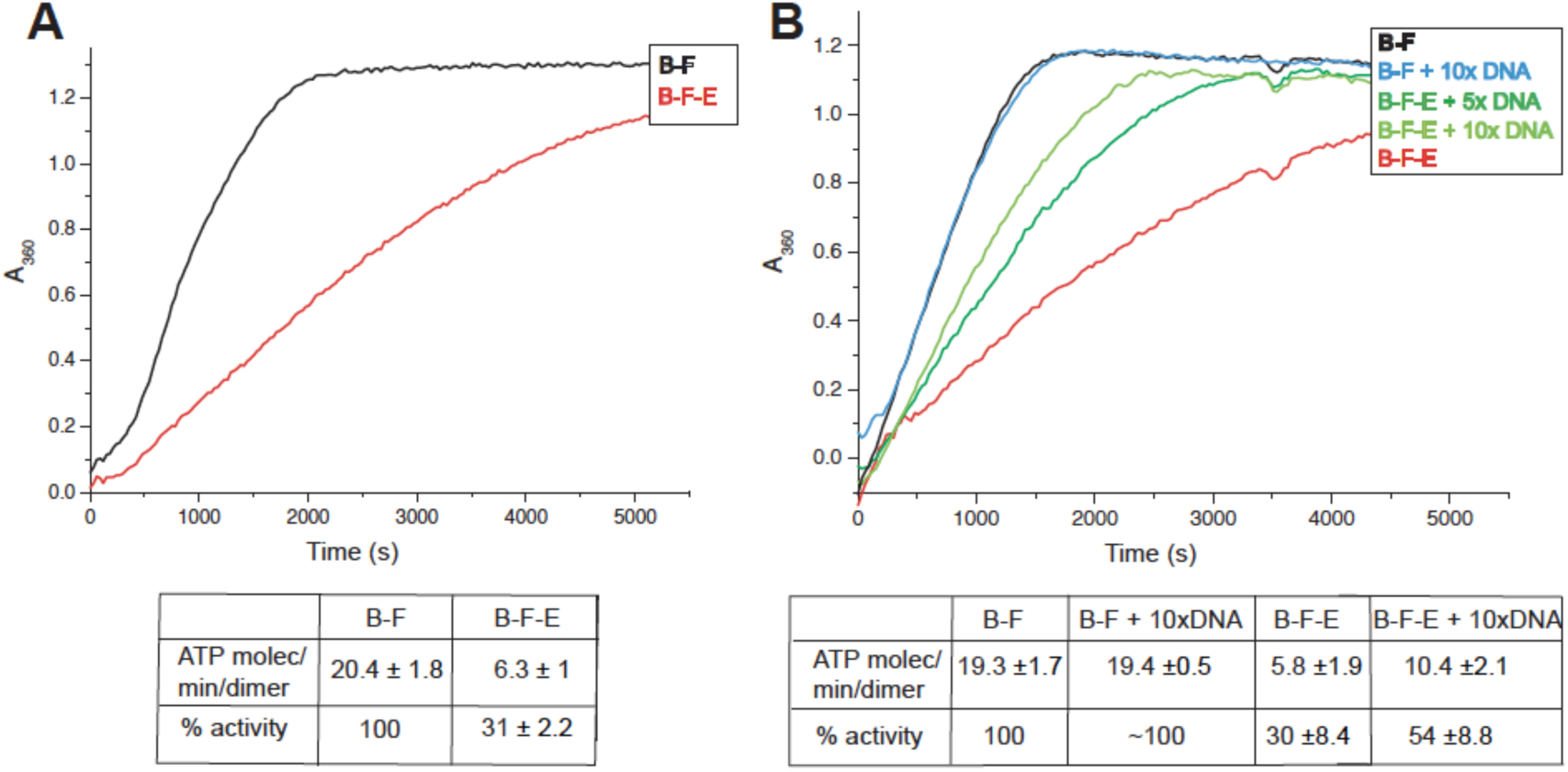
DNA relieves MukE mediated inhibition of MukF stimulated ATPase. (A) MukE– mediated inhibition. ATPase was measured at MukB, 0.5μM, and MukF 1.25μM, and MukE 5.0μM. (B) DNA alleviates MukE-mediated inhibition. ATPase was measured in the presence/absence of of 53 bp linear ds DNA fragment at 5x or 10x molar excess over MukB concentration. The average values of the initial rates +/- SD from three experiments are tabulated beneath the graphs.

### DNA binding to MukB relieves MukE-mediated ATPase inhibition

Previous reports have shown no effect of DNA on the ATPase of MukBEF (***Chen et al., 2008***, Petrushenko et al., 2006; Woo et al., 2009; ***Nolivos and Sherratt, 2014***), whereas *B. subtilis* SMC ATPase was reported to be stimulated modestly by DNA (***Hirano and Hirano, 2004***). We confirmed that MukB ATPase is independent of the presence of DNA (Figure 6B); addition of 53bp ds linear DNA at 10-fold excess (5μM) over MukB (0.5μM), did not influence MukBF ATPase activity. MukBF ATPase was not dependent on residual DNA contamination of the proteins as judged by the observation that extensive DNase treatment of MukBF did not influence the ATPase level.

DNA alleviated the MukE-mediated inhibition of MukB ATPase. The extent of this ATPase restoration was DNA concentration-dependent (Figure 6B) and at 10-fold excess of DNA over MukB, the ATPase level was restored to ∼50% of the level in the absence of MukE (Figure 7C). A similar restoration of activity was observed for most other MukF variants, although FC4 exhibited similar MukE-inhibited ATPase activities in the presence and absence of DNA (Figure 7C). MukF derivatives, FN2 and FC5, lacking MukE binding sites, and FC3, containing only the E2 binding site, were not inhibited by MukE and did not respond to DNA.

**Figure 7.**
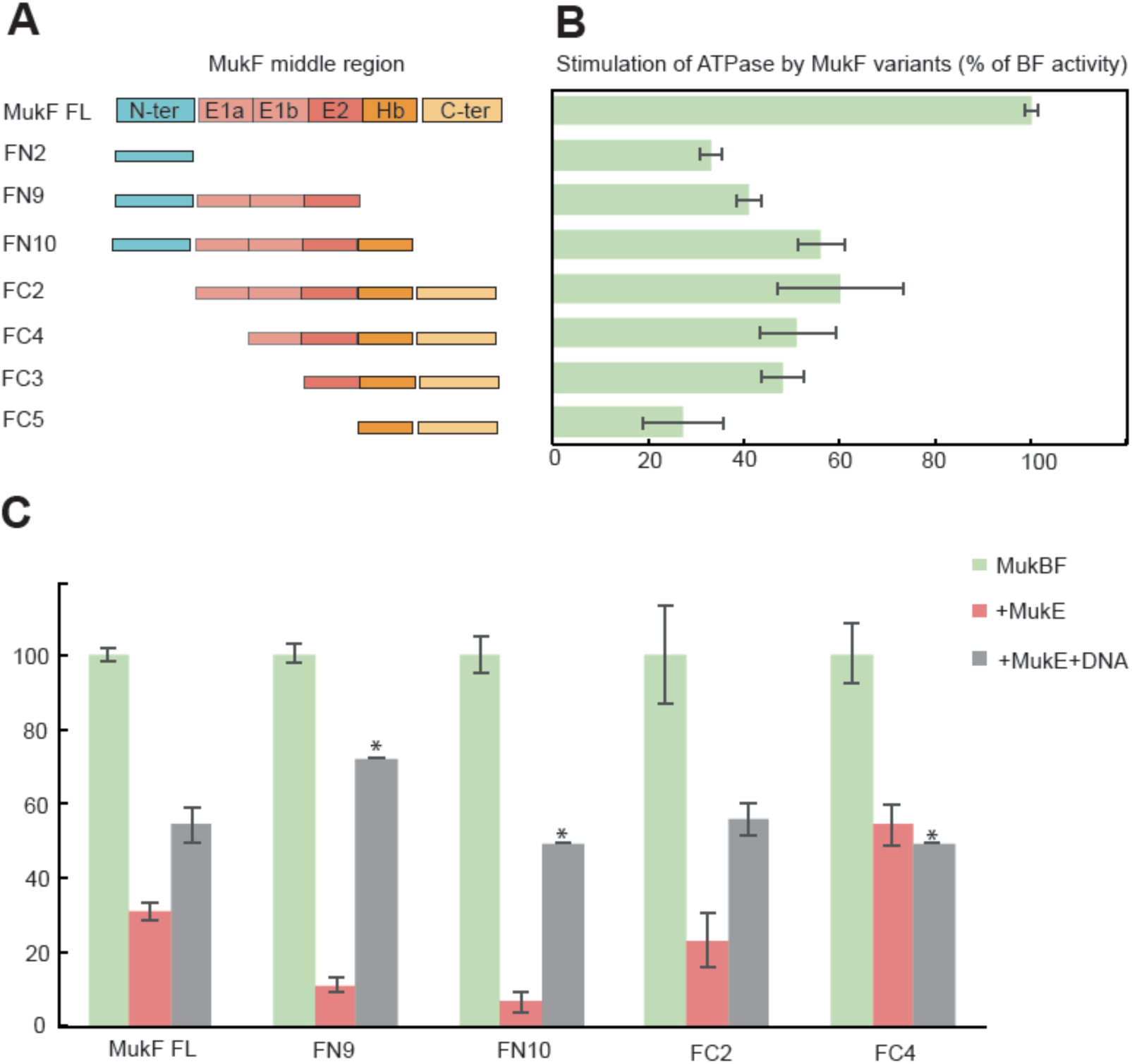
MukF middle region is important for MukF C- and N-terminal domains ATPase stimulation, and essential for MukE inhibition. (A) The extent of MukF middle region in MukF variants. As defined by the published crystal structures (Woo et al., 2009), the middle region contains two binding sites for the monomers of MukE dimer, E1 and E2. Its C-terminal part forms extended polypeptide that binds the MukB head in the asymmetric complex, Hb (Woo at al., 2009). One monomer of the MukE dimer binds helical E1a and part of the acidic linker E1b, while the second MukE monomer binds E2, also part of the extended region of the linker. Consistently with this, MukE binds to FC2, which contains entire middle region, while it doesn’t bind to FN2, which lacks the middle region (Fig. 7 supplementary figure 1). (B) Stimulation of MukB ATPase by MukF variants expressed as a percentage of stimulation by full length MukF. The bars show averages of the initial rates +/- SD from three independent experiments. (C) Inhibition of the stimulatory effect of MukF variants by MukE and its release by DNA. The MukB ATPase activated by the variants in the presence of MukE is compared to the activity in the absence of MukE as shown in B, which is considered here to be 100%. MukE did not inhibit stimulation by FN2, FC3 and FC5 variants. Release of MukE inhibition was measured in the presence of 53nt ds DNA fragment at 10-fold excess over the concentration of MukB. Only two measurements (*) were made for FN9 (range 66-77%), FN10 (range 48-49%), and FC4 (range 45-53%).

The position of the DNA binding interface on MukB heads, defined by structure-informed mutational analysis (Woo et al., 2009) indicated that DNA binding to this interface could lead to a clash with a MukE dimer bound to the MukF middle region in a heads-engaged MukBEF complex. Therefore, it seems probable that relief of MukE inhibition by DNA might reflect a competition between MukE and DNA for binding to the MukBF head complex.

### The MukF N-terminal and C-terminal domains independently modulate MukBEF action *in vivo*

Since MukF is required for MukBEF function *in vivo*, we surmised that the regulated over-expression of either the MukF C- or N-terminal domains *in vivo* might inhibit MukBEF action as a consequence of the fragments competing with endogenous MukF. MukBEF function was assessed by the presence of fluorescent MukBmYPetEF clusters observed as fluorescent foci associated with the replication origin (*ori*). MukF fragments were over-expressed, using arabinose induction, from a multicopy plasmid in cells expressing chromosomal MukBmYPetEF.

Induced over-expression of FN2 led to a rapid loss of MukBEF foci (half-life of loss ∼10 min), whereas FC2 over-expression had a much milder effect on focus loss (half-life of loss ∼45 min (Figure 8, Figure 8-figure supplement 1). In both cases, residual MukBEF clusters remained *ori*-associated. We then tested if normal cycles of ATP binding and hydrolysis are responsible for the inferred turnover of MukF within functional MukBEF complexes, by testing the effect of fragment production on MukB^EQ^EF complexes that are impaired in ATP hydrolysis and form clusters that turn over very slowly at the replication terminus (*ter*) rather than at *ori* (Badrinarayanan et al., 2012). Over-expression of either FN2 or FC2 had little effect on *ter*-associated fluorescent MukB^EQ^mYPetEF clusters, consistent with the observation in FRAP experiments that there was little turnover of these complexes, presumably as a consequence of their impaired ATP hydrolysis (Badrinarayanan et al., 2012). These observations are consistent with the hypothesis that the MukF interaction with the MukB neck turns over during cycles of ATP binding and hydrolysis, and that impairment of the interaction between MukF and the MukB neck leads to loss of functional MukBEF clusters from the chromosome. The relatively low turnover of this interaction as compared to the dwell time of MukBEF complexes *in vivo* (∼50s) and the rates of ATPase measured *in vitro* may be a consequence of the chelate effect arising from the fact that when the N-terminal domain is released from the MukB neck the reminder of MukF remains associated with the complex through interactions of its C-terminal domain and linker with MukB, thereby giving a high re-binding rate in contrast to the competing fragments in solution.

**Figure 8.**
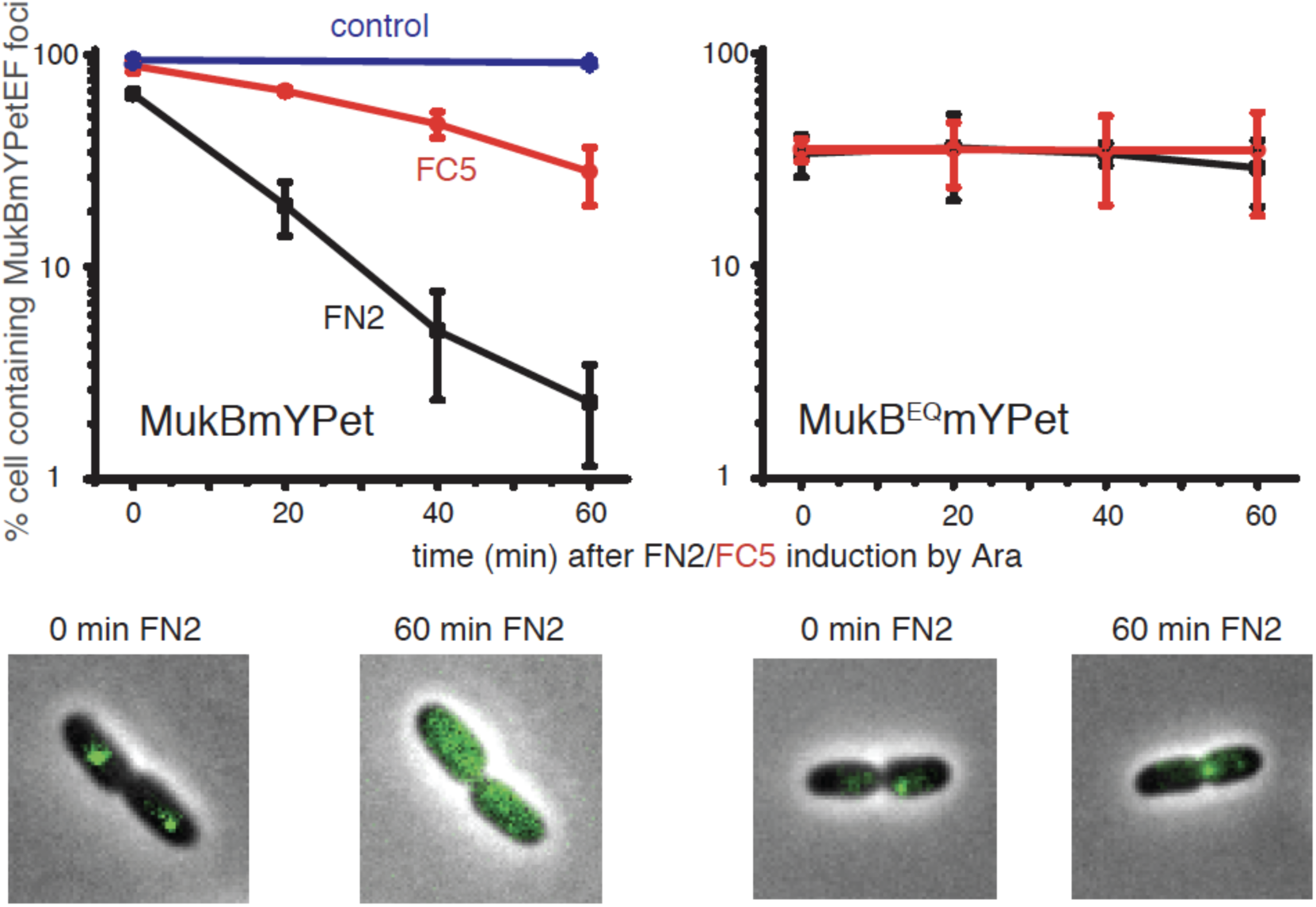
Overexpression of MukF N-terminal and C-terminal domains fragments leads to an ATP hydrolysis cycle dependent release of DNA from MukBEF complex *in vivo*. MukF FN2 and FC5 fragments were overexpressed from Para promoter in pBAD24 by addition of arabinose. MukBmYPetEF and MukB^EQ^mYPetEF complexes were visualized in the absence of arabinose and at every 20 min post induction. More than 500 cells were analysed for each condition. Experiment was repeated 3 times; error bars show standard deviation of 3 repeats. Bottom: images of FN2 overexpressing cells taken at time 0 and 60 min in MukBmYPetEF and MukB^EQ^YPetEF cells.

FN2 over-expression led to a ∼4-fold more efficient displacement of labelled MukBEF complexes from DNA, than over-expression of FC2. These results are consistent with an idea that the interface between the MukF N-terminal domain (helices 8 and 9) and the neck disengages more frequently than the interface between MukF C-terminal domain and the cap during the activity cycles of MukBEF. Indeed, the observed less frequent disengagement of the C-terminal MukF domain could arise indirectly as a consequence of turnover of the N-terminal domain. Interfaces between the kleisin N-terminus and the SMC neck in yeast, *drosophila* and human cohesin complexes have been proposed previously to function as DNA exit gates (***Beckouët et al., 2016***; Buheitel and Stemmann, 2013; Chan et al., 2012; Eichinger et al., 2013; Huis in‘t Veld et al., 2014).

## Discussion

### The functional interaction between the MukF N-terminal domain and the MukB neck breaks and reforms during cycles of ATP binding and hydrolysis

Our demonstration that the MukF N-terminal domain interacts with the MukB neck, thereby contributing to the formation of MukB(E)F tripartite rings, mirrors those observed for other SMC complexes. Indeed, mutational analysis highlights the likely structural similarity of the interface between MukF helices 8 and 9 and their potential partner coiled-coil helices in the MukB neck to the equivalent interactions in other SMC complexes (Figure 5B). The asymmetry directed by this kleisin interaction with SMC dimers therefore appears to be functionally conserved in all SMC members, regardless of whether they form homodimers or heterodimers (Bürmann et al., 2013; Gligoris, 2014; Huis in‘t Veld et al. 2014). However, unlike other kleisins, which are apparently monomeric, MukF is a stable dimer and its dimerization domain is adjacent to the 4-helix bundle, to which helices 8 and 9 belong. Nevertheless, binding of these helices to the MukB neck did not interfere with MukF dimerization, a result consistent with our previous analysis inferring the existence of MukBEF dimers of dimers *in vivo* (Badrinarayanan et al., 2012a).

The analyses reported here demonstrate that the interaction between the MukF N-terminal domain and MukB neck is not only essential for MukBEF function *in vivo*, but that this interaction is broken and reformed during cycles of ATP binding and hydrolysis. Furthermore, the observation that impairment of the normal MukF-MukB neck interactions leads to loss of MukBEF clusters from chromosomes indicates that by opening of the MukB neck-MukF interface, DNA can be released from the ‘bottom ring chamber’ formed by a kleisin bridging a MukB head and the MukB neck of a partner molecule. This result provides further support for a mechanism in which ATP hydrolysis is required to release MukBEF complexes from chromosomes (Nolivos et al., 2016). Equivalent interfaces between the kleisin and SMC3 neck in the yeast, *drosophila* and human cohesin complexes have also been proposed to act as DNA exit gates and it has been proposed that this interaction, which is not required for loading onto chromosomes, turns-over in response to ATP binding and hydrolysis (Beckouët et al, 2016; Buheitel and Stemmann, 2013; Chan et al., 2012; Eichinger et al., 2013; ***Eltbash et al., 2016).***

Although, a DNA exit gate formed by the SMC coiled-coil neck-kleisin interaction appears to be conserved, we think it possible that other interfaces could also be used for DNA release under certain circumstances. For example, the hinge dimerization interface, which has been proposed to be a DNA entrance gate (Buheitel and Stemmann, 2013; Gruber at al., 2006), might additionally function as an exit gate under some conditions (Murayama and Uhlmann, 2013; Uhlmann 2016). Because there are two potential proteinaceous chambers in SMC complexes, the upper one formed by a heads-engaged SMC complex and the lower one by the kleisin bound to the SMC (Diebold-Durand et al., 2017; Uhlmann, 2016;), then each of these chambers could have exit (and entrance) gates for DNA segments entrapped within each of them. In MukBEF, interaction of MatP-*matS* with the MukB hinge has been proposed to promote ATP hydrolysis-dependent release of MukBEF clusters from the *ter* region of the chromosome, suggestive of release through the dimerization hinge (Nolivos et al., 2016). Similarly, MukB-dependent stimulation of catalysis by TopoIV could arise as a consequence of DNA exiting the MukB hinge and being presented to the TopoIV entrance gate, which is in proximity to the MukB hinge (Vos et al., 2013).

### Two independently regulated ATPases in asymmetric MukBF complexes

The work reported here provides new insight into how kleisin interaction with the SMC heads leads to an asymmetric complex that has two putative independently controlled ATPases, one activated by the kleisin C-terminal domain and the other by the N-terminal domain. Since we have inferred previously from *in vivo* experiments that ATP hydrolysis by MukBEF is required for both loading and unloading onto DNA (Nolivos et al., 2016), a parsimonious explanation of our data is that the MukB neck-kleisin interaction acts in DNA unloading, leaving the MukB head-kleisin interaction to function in loading onto chromosomes. Similarly, a functional asymmetry in yeast cohesion ATPase active sites has been proposed, with the one equivalent to the kleisin-neck interaction uncovered here being required for release from chromosomes, while both ATPases were implicated in loading onto chromosomes (Çamdere et al., 2015; ***Elbash et al., 2016***). Our own data do not address whether the interaction with the MukB neck is also required for loading.

Asymmetric ATPase mechanisms have also been demonstrated for some ABC transporters, which share the overall organisation of their ATPase heads with SMCs (Beek et al., 2014; Procko et al., 2017; Zhou et al., 2016). Similarly, eukaryotic heterodimeric SMC complexes have active site region differences; for example, comparison of SMC1 sequences with those of SMC3 show protein family specific differences, with their ATPases being differentially regulated (Beckouët et al., 2016; Çamdere et al., 2015; ***Elbash et al., 2016***). It remains to be determined how the two different domains of MukF independently stimulate activity of two sister ATP active sites and how these two ATP binding and hydrolysis steps fit into the multiple stages of the MukBEF activity cycle on chromosomes.

### MukE and DNA compete during cycles of ATP binding and hydrolysis

The functional significance of how MukE binding to MukF inhibited ATPase activities stimulated by either the MukF C-terminal, or the N-terminal domains, remains unclear. It could arise simply from the fact that MukE binding stabilizes a particular MukBF conformation, thereby leading to less turnover during the steady state multiple turnover ATPase assays.

Alternatively, or additionally, this could reflect MukE playing a regulatory role during transitions between various stages of MukBEF activity cycle. Other *in vitro* and *in vivo* studies have postulated a regulatory role of MukE (Gloyd at al., 2011; She at al., 2013), although details of how this regulation is mediated have been unclear. Nevertheless, depletion of MukE *in vivo* mimics the ATP hydrolysis-impaired phenotype of a MukB^EQ^ mutant, which loads slowly onto DNA in the *ter* region but is unable to undergo the multiple cycles of ATP binding and hydrolysis required to re-locate to *ori* (Badrinarayanan et al., 2012b; Nolivos et al., 2016).

Analysis of steady state ATPase levels indicate MukE and DNA compete for binding to the MukB head. This is consistent with *in vitro* studies, which showed competition between MukEF and DNA for MukB binding and that MukEF inhibited MukB-mediated DNA condensation (Cui et al., 2008; Petrushenko et al., 2006,). Furthermore, a patch of positively charged amino acid residues on the surface of MukB head, close to the base of the neck, was shown to be important for interaction with DNA (Woo at al., 2009). Projection of B form DNA onto this patch highlights the potential competition of DNA- and MukE-binding to a MukBF complex, which may reflect alternative states during the MukBEF-DNA activity cycle.

### Perspective

A range of SMC complex structures and extensive biochemical and functional analyses lead to the conclusion that all SMC complexes, including MukBEF, share distinctive architectures and similarities in their likely molecular basic mechanisms of action on chromosomes. The asymmetric architecture that directs the asymmetric ATPases revealed here, and with other SMC complexes, along with the kleisin-SMC neck interaction being important for ATP hydrolysis-dependent release of chromosomal DNA, are two such conserved features. Central to the SMC complex mechanism is the ability to bind and hydrolyse ATP in a regulated fashion, which directs stable loading onto chromosomes, rapid transport with respect to DNA, and regulated release from chromosomes (Diebold-Durand et al., 2017; ***Terekawa et al., 2017; Wang et al., 2017***). Unlike other SMC complexes, cohesins have two distinct and separable activities. In addition to their roles in facilitating organisation of non-replicating chromosomes (***Çamdere et al., 2016***; Uhlmann, 2016), they have evolved the ability to held newly replicated sisters stably together (cohesion), until controlled release is triggered. The latter activity requires that ATPase-hydrolysis-dependent turnover of cohesion complexes on chromosomes, mediated by Wapl, is inhibited by acetylation of the SMC3 subunit, which presumably inhibits SMC3 ATPase (***Ben-Shahar et al.***, Chan et al., 2012; Kueng et al., 2006; Unal et al., 2008). In addition to this inhibition of turnover, cohesion requires entrapping of the two sister chromatids within the SMC complexes, an activity separable from that involved into loading onto single chromosomes, and a process that must involve an altered association of DNA segments within the SMC complex.

The detailed mechanism by which SMC complexes transport themselves with respect to DNA remains elusive, although recent experiments have demonstrated autonomous SMC ATP-hydrolysis-dependent DNA transport *in vitro* (***Terekawa et al., 2017***). Any such transport must require at least two specific DNA-SMC complex attachment points on different conformational states of the complex, with coordinated transitions as transport proceeds. Our demonstration that MukF dimerization is maintained during its interaction with the MukB neck, not only supports our demonstration of dimers of MukBEF dimers in active MukBEF clusters *in vivo*, but provides additional support for our previously proposed ‘rock (rope) climber’ model for the transport of MukBEF dimer of dimers with respect to DNA (Badrinarayanan et al., 2012a). This model assumes that dimers of MukBEF dimers are a minimal functional unit, in which coordinated capture and processing of DNA segments by each MukBEF dimer, similar to the action of a climber reaching out to ‘grab’ a rock/rope alternatively with each arm. The staggered cycles of ATP binding and hydrolysis, DNA trapping and release and associated with them conformational changes could effectively coordinate the activity within the partner dimers within MukBEF dimers. For SMC complexes that do not obviously form dimers of dimers, the type of model proposed by Diebold-Durand et al. (2017), in which DNA loops captured in the upper SMC chamber are transferred to the lower chamber, where they fuse with a pre-existing loop, thereby meeting the basic requirements for ATP hydrolysis-driven transport.

## Materials and Methods

### Protein Purification

MukB, MukB_H,_ MukB_HN_, MukE, were 6xHis-tagged at the C-terminus, while MukF and its C- and N-terminal truncations were 6xHis-tagged at the N-terminus and were expressed from plasmid pET21 and pET28, respectively in C3013I cells (NEB). 2L cultures were grown in LB with appropriate antibiotics at 37°C to A_600_∼0.6 and induced by adding IPTG at final concentration of 0.4 mM. After 2 hours at 30°C, cells were harvested by centrifugation, re-suspended in 30ml lysis buffer (50 mM HEPES pH 7.5, 300 mM NaCl, 5%glycerol, 10 mM imidazole) supplemented with 1 tablet of protease inhibitor (PI), and homogenized. Cell debris was removed by centrifugation and clear cell lysates were mixed with 5 ml equilibrated TALON Superflow resin, poured into a column, then washed with 10 X volume of washing buffer (50 mM HEPES pH 7.5, 300 mM NaCl, 5% glycerol, 25 mM imidazole, PI). Bound proteins were eluted in elution buffer (50 mM HEPES pH 7.5, 300 mM NaCl, 5% glycerol, 250 mM imidazole). The fractions from TALON were diluted to 100 mM NaCl buffer and injected to HiTrapTM Heparin HP column (GE Healthcare) pre-equilibrated with Buffer A (50 mM HEPES pH 7.5, 100 mM NaCl, 10% glycerol, 1 mM EDTA, 1 mM DTT), then the column was washed at 1 ml/min flow rate until constant UV280. Purified fractions were eluted with a gradient 100-1000 mM NaCl.

For MukE and MukF purifications, fractions from Talon were diluted and injected to HiTrap DEAE FF column (GE healthcare) pre-equilibrated in Buffer A. Purified fractions were eluted with a gradient 100 -1000 mM NaCl. Protein concentration was estimated by UV absorption at 280 nm on Nanodrop spectrophotometer, and protein purity and identity confirmed by electrospray ionization mass-spectrometry and SDS PAGE. Proteins were aliquoted and stored at -20°C in a buffer containing 10% glycerol.

### ATP Hydrolysis Assays

ATP hydrolysis was analysed in steady state reactions using an ENZCheck Phosphate Assay Kit (Life Technologies). 150 μL samples containing standard reaction buffer supplemented with 2 mM of ATP were assayed in a BMG Labtech PherAstar FS plate reader at 25° C. The results were computed using MARS data analysis software. Quantitation of phosphate release was determined using the extinction coefficient of 11,200 M^-1^cm^-1^ for the phosphate-dependent reaction at 360 nm at pH 7.0.

### Size Exclusion Chromatography and Multi-Angle Light Scattering (SEC-MALS)

Purified proteins were fractionated on a Superose 6 10/300 GL or a Superose 12 10/300 column equilibrated with 50 mM HEPES, pH 7.5 buffer containing 100 mM NaCl, 1 mMDTT, 1 mM EDTA, at flow rate of 0.5 ml/min. 500 μl samples containing analysed proteins were injected on the column and run at a flow rate of 0.5 ml/min. SEC-MALS analysis was performed at 20°C using a Shimadzu (Kyoto, Japan) chromatography system, connected in-line to a Heleos8+multi-angle light scattering detector and an Optilab T-rEX refractive index (RI) detector (Wyatt Technologies, Goleta, CA). Protein samples in 50 mM HEPES pH 7.5, 100 mM NaCl, 1mM DTT, 1mM EDTA, 10% glycerol, were injected in this system, and the resulting MALS, RI and UV traces processed in ASTRA 6 (Wyatt Technologies).

### Pull-down Assays

MukF FLAG-tagged fragments were expressed from pET DUET plasmids in C3013I cells (NEB).1L cultures were grown in LB with carbenicilin (100 μg/ml) at 37°C to A_600_∼0.6 and induced by adding IPTG to a final concentration 0.4 mM. After 2 hours at 30°C, cells were harvested by centrifugation, re-suspended in 30ml lysis buffer (50 mM HEPES pH 7.5, 300 mM NaCl, 5%glycerol, 10 mM imidazole) supplemented with 1 tablet of protease inhibitor (PI), and homogenized. Cell debris was removed by centrifugation and clear cell lysates were mixed with 150 μl Anti-FLAG M2 Affinity gel (Sigma Aldrich), incubated for 1 hr at 4°C. The resin was then washed 3 times with the same buffer containing 250 mM NaCl, resuspended in 1ml of buffer I (50 mM HEPES pH 7.5, 100 mM NaCl), and purified MukB, MukB_H_ or MukB_HN_ were added. After 45 min incubation (4°C) the resin was washed 3 times, re-suspended in 200 μl of protein loading buffer (NEB) and analyzed on 4-20% gradient SDS PAGE.

### Mutagenesis

Point mutations in plasmid-encoded genes were made using Q5 Site-Directed Mutagenesis Kit (NEB). Primers were designed with NEBase Changer. 10 ng of the template was taken to the reaction. Plasmids were isolated and mutations confirmed by sequencing.

### Complementation assays

The ability of leaky plasmid-encoded MukB expression from pET21, in the absence of IPTG, to complement the temperature-sensitive growth defect of **Δ***mukB* AB1157 cells at 37°C in LB was assayed. Cells were transformed with pET21 carrying MukB or MukB variants and allowed to recover for 8 hr post transformation at permissive temperature then plated in duplicates on LB plates containing carbenicillin (100μg/ml). One plate was incubated at non-permissive (37°C) and the other one at permissive (20°C) temperature. Six colonies from plates incubated at permissive temperature were streaked in duplicate and grown at permissive and non-permissive temperature along with positive and negative controls.

### Analysis of MukBEF function *in vivo*

Strains were streaked onto LB plates with appropriate antibiotics. Single colonies were inoculated into M9 glycerol (0.2%) and grown overnight at 37°C to A_600_ 0.4-0.6, then diluted into fresh M9 and grown to A_600_ 0.1. Cells were spun and immobilized on agarose pads between two glass coverslips (1.5 thickness). 1% agarose pads were prepared by mixing low-fluorescence 2% agarose (Bio-Rad) in dH_2_O 1:1 with 2x growth medium. For analysis of MukBEF fluorescent clusters (foci), strains carrying either functional MukBmYPet (SN182), or the ATP hydrolysis-impaired mutant MukB^EQ^mYPet (SN311, Nolivos et al., 2016) were used. Wide-field fluorescence microscopy used an Eclipse TE2000-U microscope (Nikon), equipped with an 100x/NA1.4 oil PlanApo objective and a Cool-Snap HQ^2^ CCD, and using Metamorph software for image acquisition. Over-expression of FN2 and FC2 was from pBAD24 plasmids containing the appropriate arabinose-inducible MukF derivative. Strains were transformed with given plasmid and grown in M9 glycerol medium supplemented with 0.2% glucose to limit leaky expression from the arabinose promoter. Once cultures reached A_600_ ∼0.1, cells were centrifuged and re-suspended in M9 glycerol medium supplemented with 0.2% L-Arabinose and grown at 37°C. Every 20 minutes, cells from1 ml of culture were taken, centrifuged, placed on agarose pad and imaged. As a control, strain carrying empty pBAD24 vector was analysed. Cells were segmented from brightfield images using MicrobeTracker (Sliusarenko et al., 2011). MukB foci were detected using ‘spotfinderM’, available as part of the MicrobeTracker Suite.

### Fluorescence Correlation Spectroscopy (FCS)

FCS was carried out on a ConfoCor 2 system (Carl Zeiss). The 633nm line of a HeNe laser was directed via a 488/561/633 dichroic mirror and focused with a Zeiss C-Apochromat 40Å∼ NA 1.2 water immersion objective to excite experimental samples containing Cy5. Fluorescence emission was collected using a 655-nm long pass filter and recorded by an avalanche photodiode. The pinhole diameter was adjusted to 83μm (one Airy unit), and the pinhole position was optimized with use of the automatic pinhole adjustment for Cy5. All FCS experiments were carried out in Lab-Tek (Nagle Nunc International) eight-well chambered borosilicate glass plates at 22 ± 1 °C. In the assay, diffusion of Cy5-labelled FN2 and FN10 fragments at fixed concentrations (∼10nM) was measured in samples carrying MukB at a range of concentrations up to 160μM. Since MukB is much larger than any of the fragments used, up to a 3-fold increase in diffusion time was observed.

The intensity of fluorescence signal was measured and the autocorrelation function G(t) was determined for diffusing fluorescently labelled species present in the sample. If two species with different diffusional properties are present, the autocorrelation function G(t) can be described as a two-component model that allows analysis of the abundance of each species:

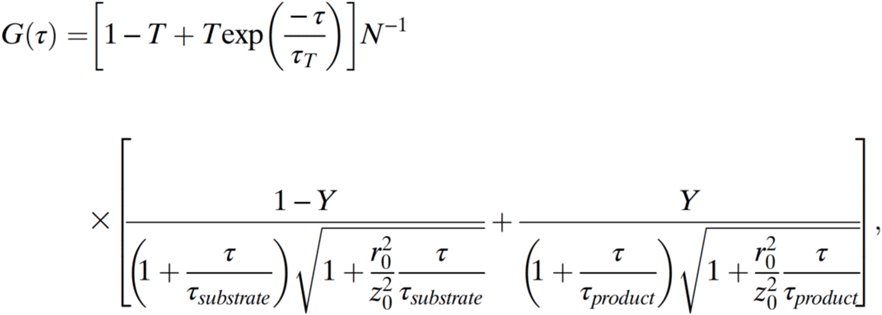

where T is the average fraction of dye molecules in the triplet state with the relaxation time **τ**T, N is the average number of fluorescent molecules in the volume observed, Y is the relative fraction of fragment bound to MukB, **τ** substrate and **τ** product are the diffusion time constants of free protein (labelled fragment as indicated for individual experiment and fragment bound to MukB), respectively, and r0 and z0 are the lateral and axial dimensions, respectively, of the observation volume. All calculations, including the evaluation of the autocorrelation curves, which was carried out with a Marquardt nonlinear least-square fitting procedure, were performed using the ConfoCor 2 instrument software. To obtain the % of bound and unbound fragments, the diffusion times for fluorescently labelled fragment were measured and fixed during data analysis. The diffusion time for the complex of a given fragment and MukB was estimated based on measured diffusion time for labelled MukB. No change in diffusion time for labelled MukB was observed when unlabelled fragment was added; therefore, the measured diffusion time for MukB was used as a fixed value during data analysis.

### Fluorescence polarization anisotropy (FPA)

Experiments were done on BMG LABTECH PHERAstar FS next-generation microplate reader with an FP 590-50 675-50 optic module. Samples were measured in Corning black 96 well flat bottom half volume plates at 25 °C. All sample volumes were 100 uL. Cy5 labelled FN3 and FC2 were used at 5 nM and 9 nM respectively. The concentration of MukB was varied from 0.1 nM to 1 μM. Samples were equilibrated for 40 minutes before measurement. Experiments were repeated thrice and standard deviations are reported. Data were plotted and analysed using Sigmaplot, where *K*_*d*_ and total receptor concentration were solved simultaneously. Binding reached saturation above 160 nM MukB. Binding of FN3 or FC2 with 1 μM MukB was used as a 100 % bound reading. Fraction of FN3 or FC2 bound was determined by using the equation:

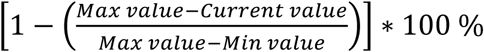

Data were plotted and the value of *K*_*d*_ and ‘total receptor’ concentration (R_T_) were simultaneously determined using Sigmaplot by solving the quadratic for fraction bound (B) below,

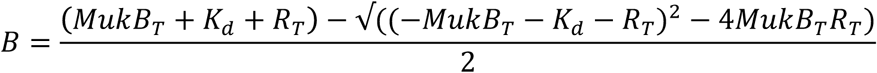

## Acknowledgements

The work was supported by a Wellcome Trust Senior Investigator Award to DJS (099204/Z/12Z) and by the Leverhulme Trust (RP2013-K-017). We thank all past and present DJS laboratory members for illuminating discussions.

## Contributions

KZ carried out most of the experimental work, including construction of all protein variants, pull-down assays, SEC-MALS, ATPase, and FCS assays. PZ performed *in vivo* analyses and contributed to FCS analysis. KR carried out FPA assays and structure modelling. RB carried out the SEC analyses, ATPase and *in vivo* complementation assays. All authors contributed to data interpretation and presentation. LKA and DJS designed and supervised the project and wrote the manuscript.

## Supplementary Figures

**Figure 3 – figure supplement 1.**
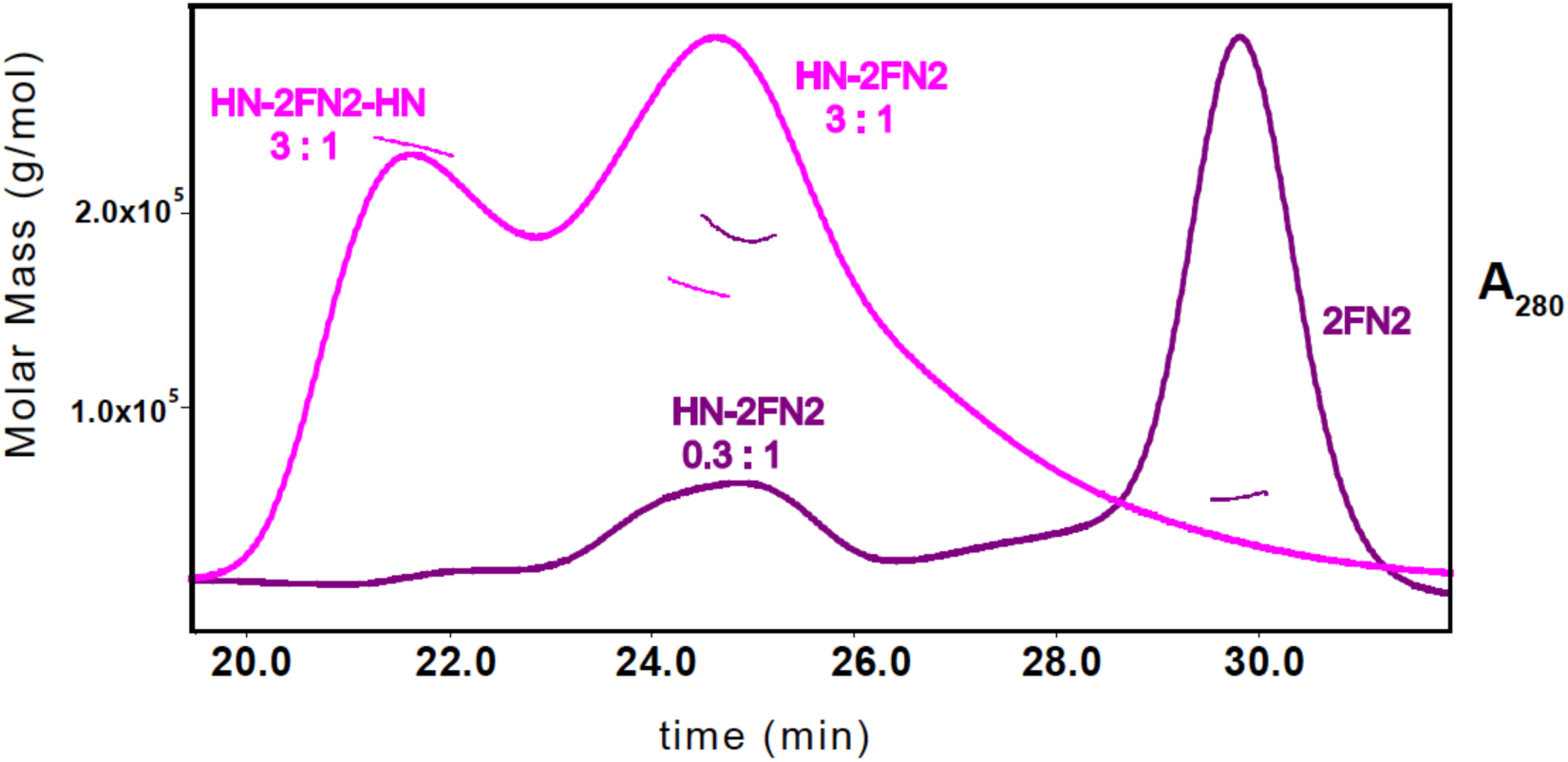
SEC-MALS analysis of MukB_HN_-2FN2 complexes. The samples contained a mixture of MukB_HN_ and FN2 at two different ratios: 3:1 m:d, pink; and 0.3:1 m:d, purple.

**Figure 3 – figure supplement 2.**
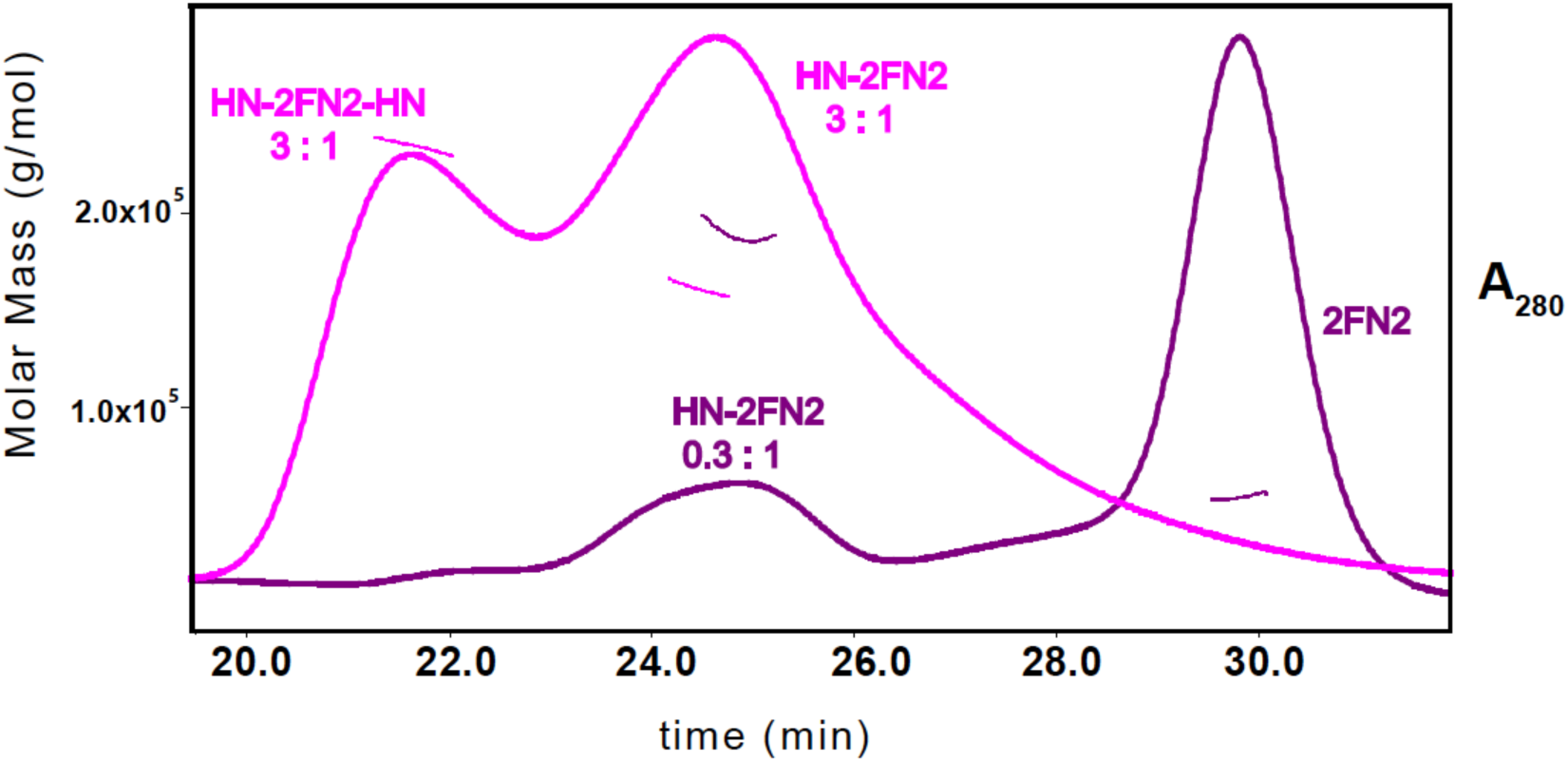
SEC-MALS analysis of MukB_HN_–2FN2 and MukB_HN_–FN2–FC2 complexes in the absence and presence of ATP (1mM).

**Figure 3 – figure supplement 3.**
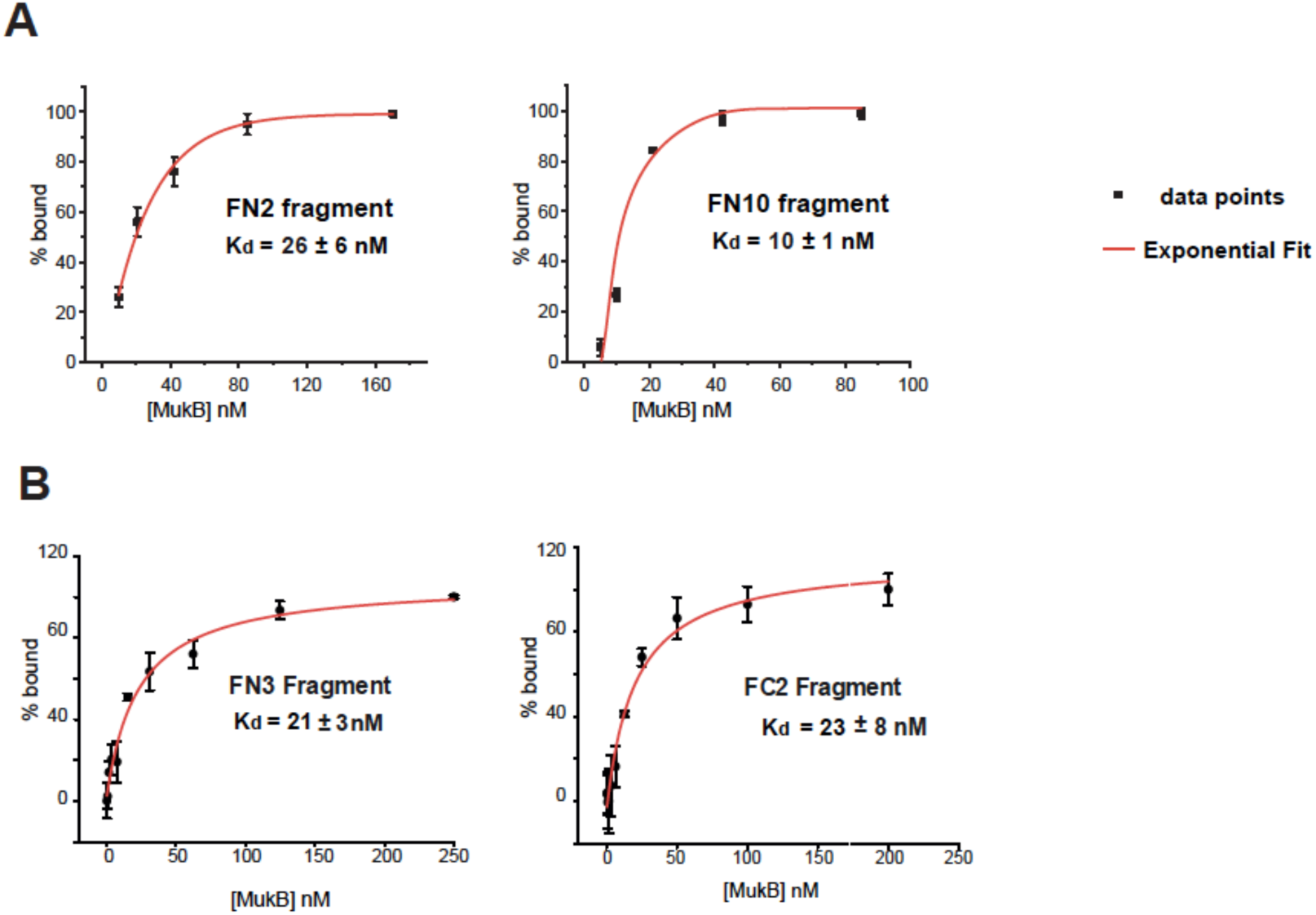
Binding affinities of MukF fragments to MukB (A) FCS measurements of FN2 and FN10 binding to MukB. Cy5 labelled fragments were at a fixed ∼10nM concentration; exponential fit to data points was used to extract Kd. Error bars represent S.D. of three independent experiments (B) FPA measurements of FN3 and FC2 binding to MukB. Cy5 labelled FN3 and FC2 fragments were at concentration 5nM and 9nM, respectively.

**Figure 4 – figure supplement 1.**
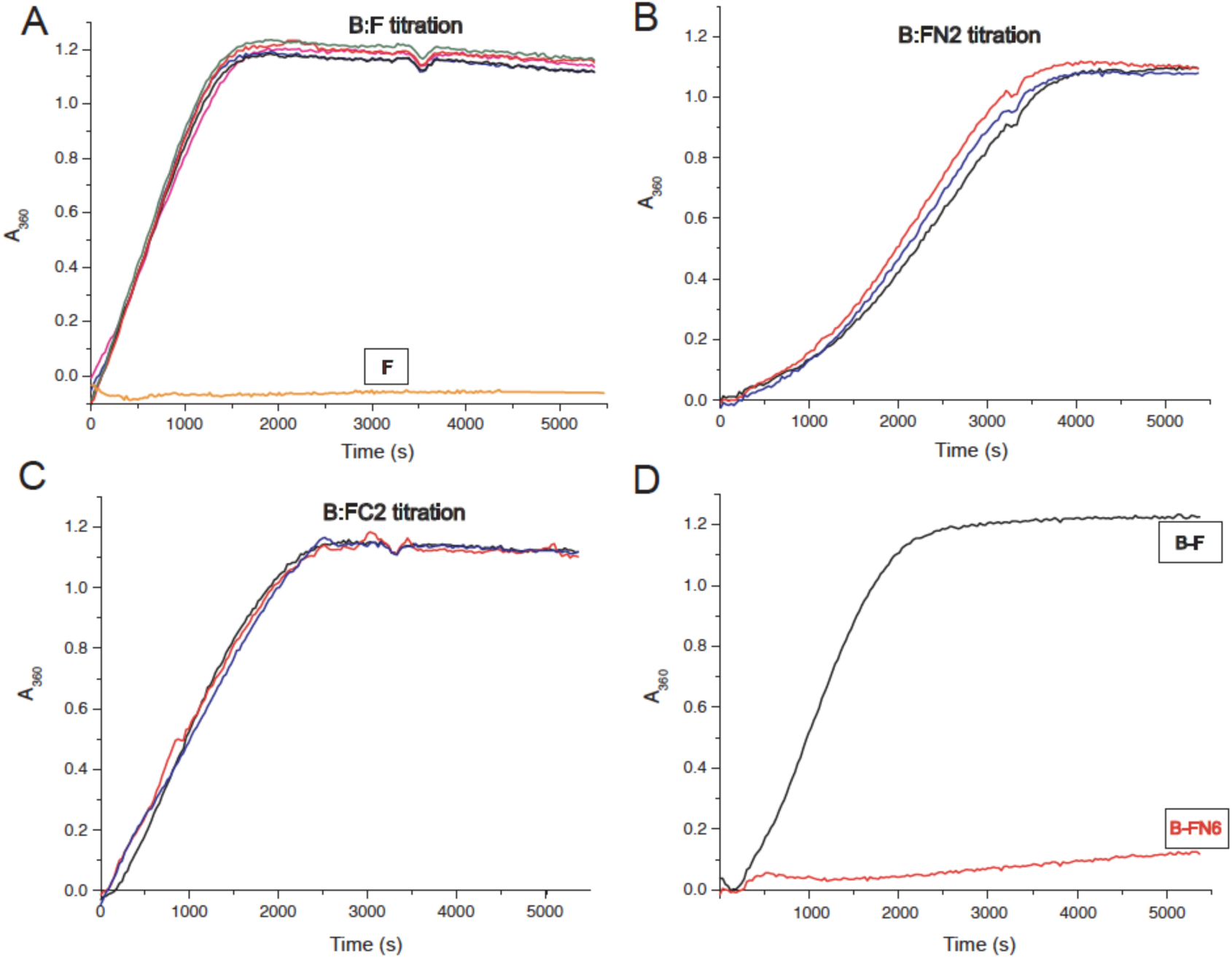
MukF stimulated MukB ATPase. (ABC) MukB ATPase activity at a range of concentrations of MukF/FN2/FC2. MukB was present at 0.5μM, MukF/FN2/FC2 at following concentrations: 0.25μM pink, 0.5μM black, 1.25μM blue, 2.5μM red, and 5μM green (in B and C 0.6μM black). (D) MukF FN6 failed to stimulate MukB ATPase. (E) MukB_HN_ ATPase in the presence of MukF, FN2 and FC2. The traces represent values from one of three replicates of the experiment.

**Figure 5 – figure supplement 1.**
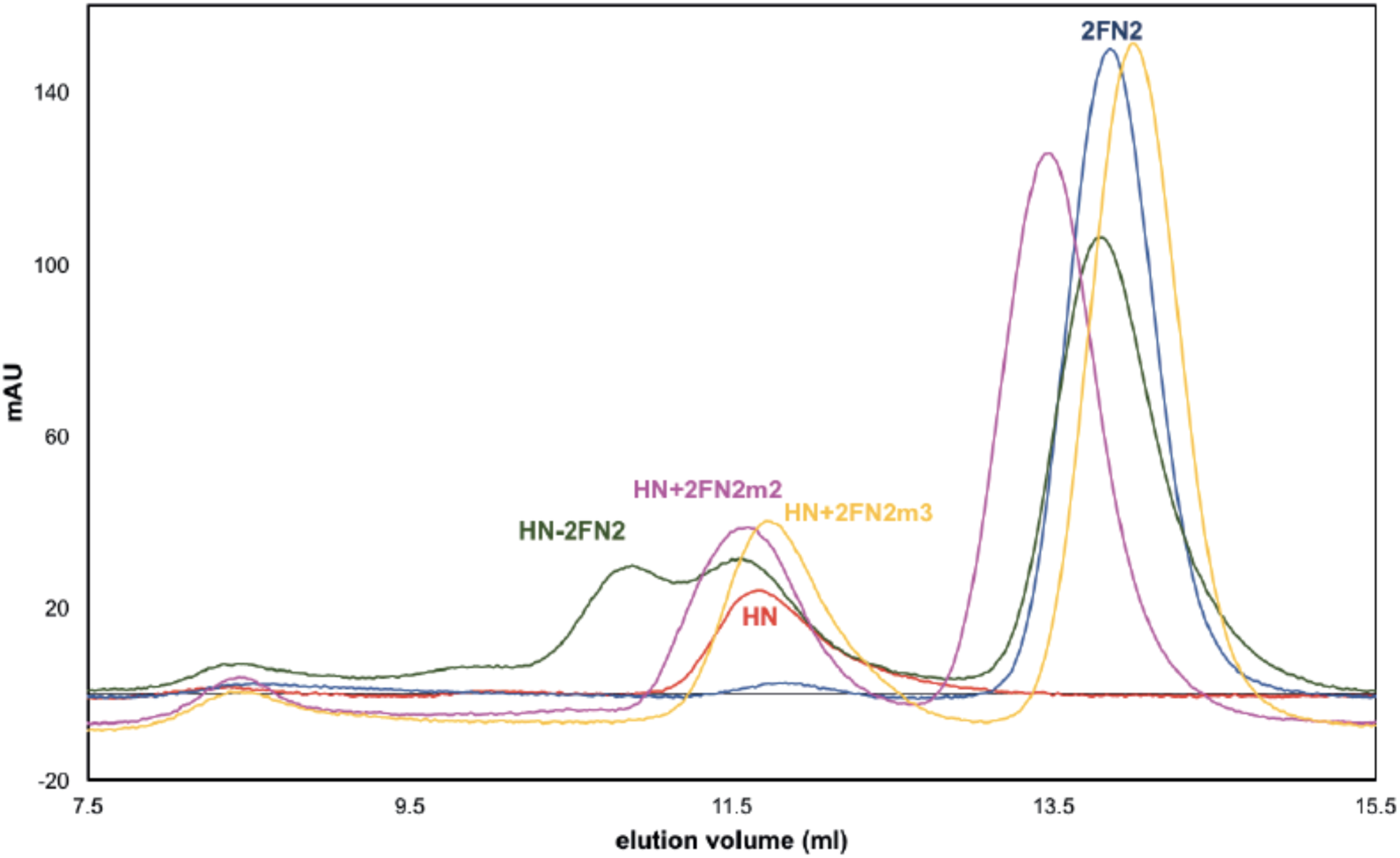
Mutated FN2 fragments were defective in binding to MukB_HN_. Binding of FN2 (dark grey trace), FN2m2 (R279E K283A R286A, pink trace), and FN2m3 (D261K S265K Q268A, yellow trace) to MukB_HN_ was analyzed by SEC on Sephadex 200 column using MukB_HN_ alone (red trace) and FN2 alone (navy blue trace) as reference. The protein concentrations were: MukB_HN_ - 3.7μM (monomer) and FN2 4.6 μM (dimer).

**Figure 5 – figure supplement 2.**
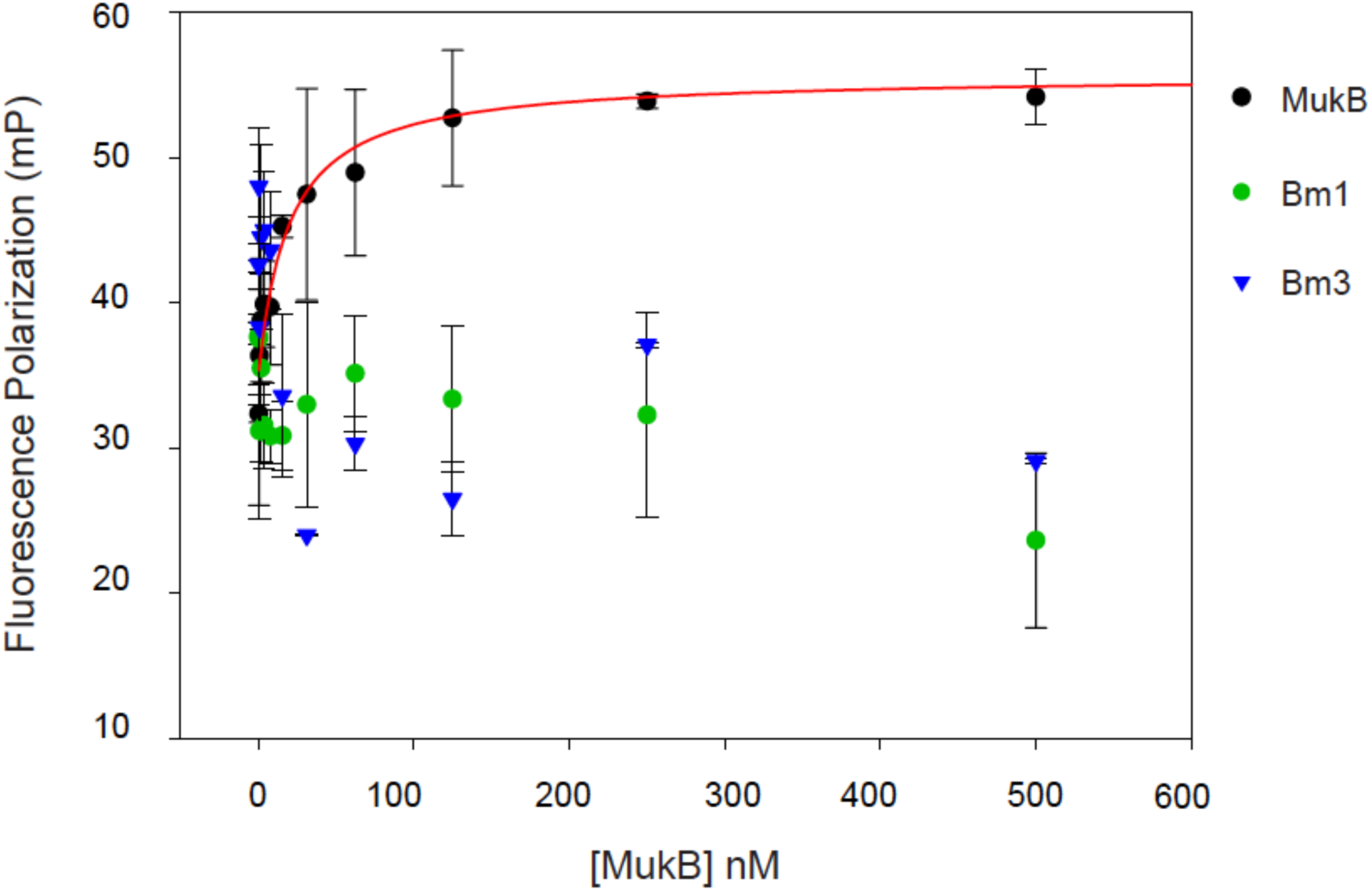
Mutated MukBm1 and MukBm3 fail to bind MukF N-terminal fragment. Binding of Cy5-labelled FN3 (at concentration of 5nM) was assessed by FPA.

**Figure 5 – figure supplement 3.**
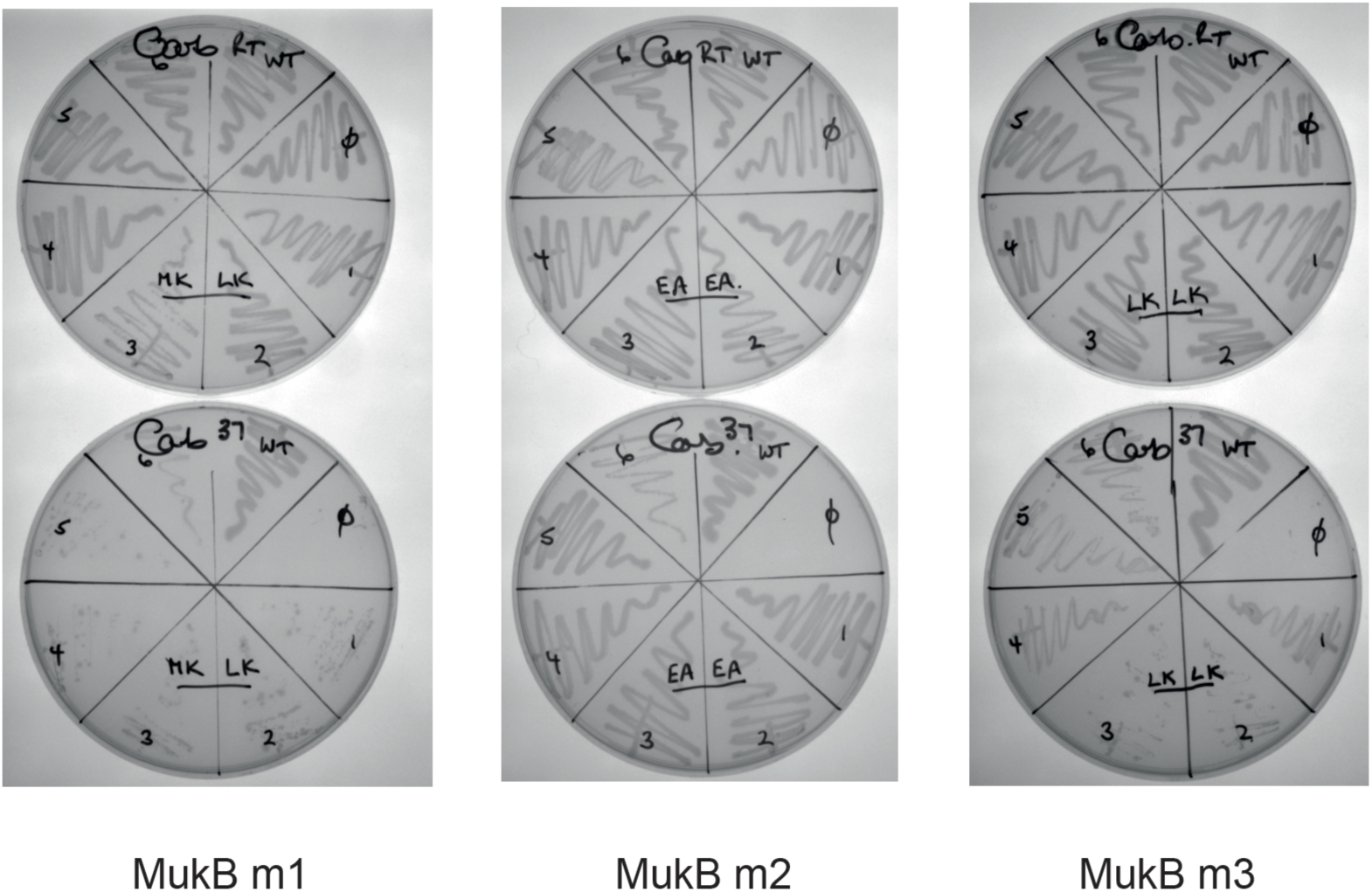
Functional analysis of mutated MukB neck variants. *in vivo* complementation in cells lacking chromosomal mukB gene by variants expressed from pET21 plasmid was assessed in the absence of IPTG (constitutive leaky expression). Growth of material streaked from 6 colonies of each variant was compared to growth of cells carrying WT MukB construct at permissive, 20°C, and non-permissive, 37°C; ϕ - a negative control, empty vector.

**Figure 5 – figure supplement 4.**
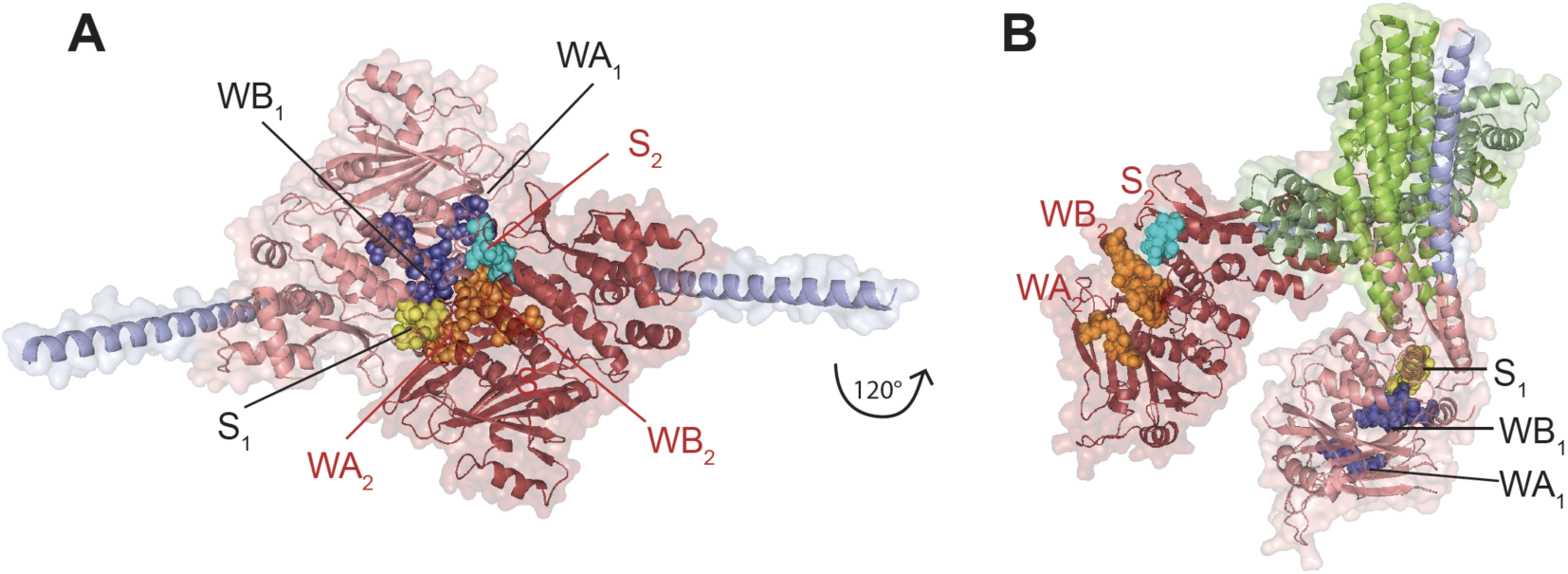
Model of the complex made by FN2 dimer binding two monomers of MukB_HN_. (A) Cartoon of the model of MukB_HN_ complex based on the asymmetric complex structure from ***Woo et., al 2009***, as shown in Figure 1B, but viewed from the top. MukB_HN_, monomer 1, is coloured salmon pink, monomer 2, intense red. The residues that make the motifs of ATPase catalytic binding sites are shown as spheres and indicated as follows: in monomer 1: Walker A and Walker B are coloured blue and signature loop in yellow, while in monomer 2: Walker A and Walker B are orange and the signature loop is in cyan; the modelled helices of the necks are shown in lilac. In this conformation two assembled active sites, WA1+WB1+S2 and WA2+WB2+S1, bind two nucleotide molecules (not shown here). (B), A model of the 2FN2-2xMukB_HN_ complex inferred from the studies presented here; the model assumes a fixed conformation of FN2 dimer. Interactions between the two independently bound MukB_HN_ necks and the helices of four-helix bundle of MukF N-terminal domain impose a conformation, in which the heads are turned around with respect to one another separating the catalytic active sites’ motifs. MukB_HN_ monomer 2 is shown in the ‘fixed’ position in both panels, although it is rotated by 120° to allow visualization of the difference of the relative arrangement of head monomers.

**Figure 6 – figure supplement 1.**
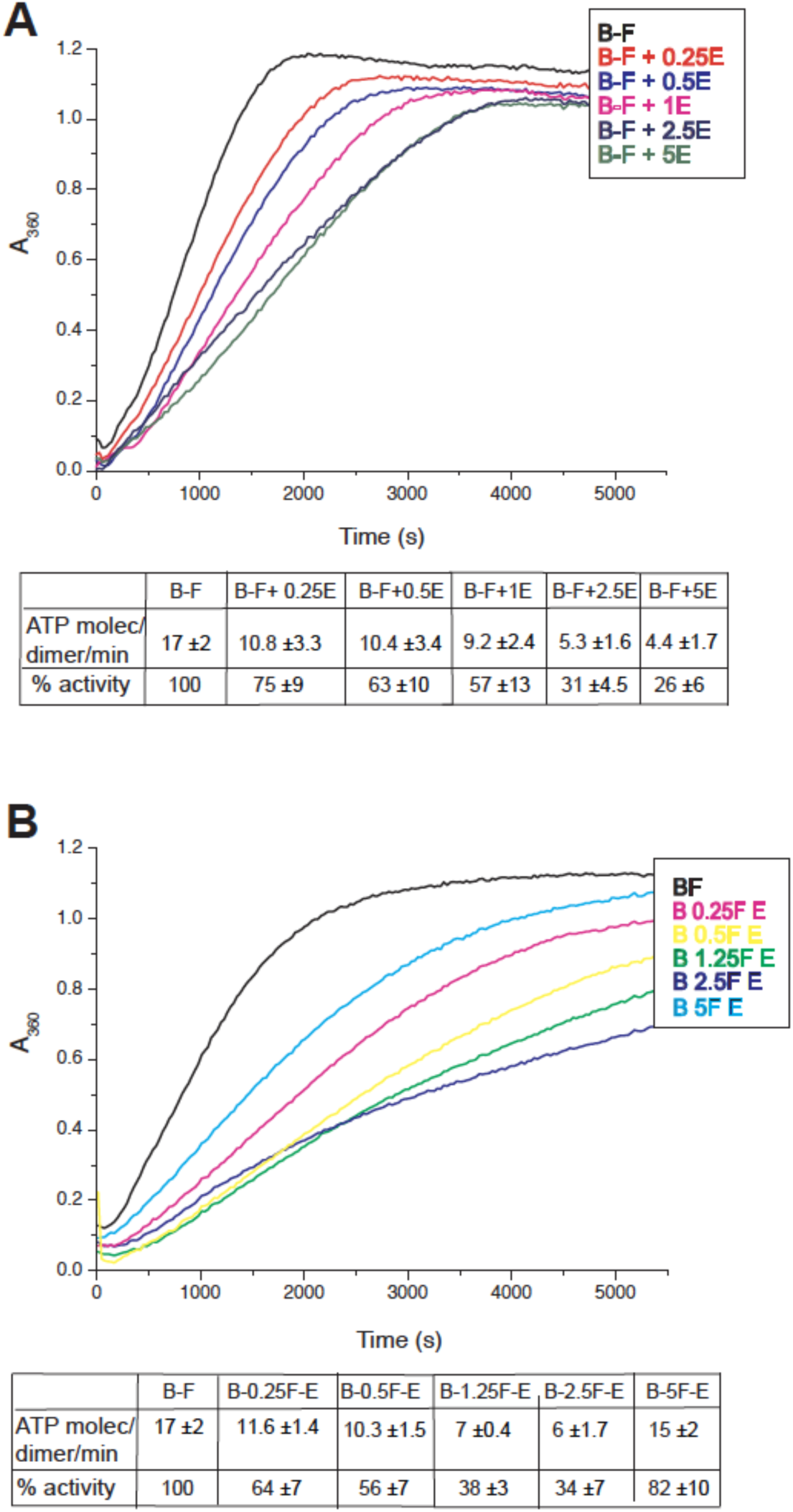
Inhibition of MukBF ATPase by MukE. (A) ATPase activity was measured at constant concentrations of MukB, 0.5μM and MukF 1.25μM and a range of concentrations of MukE from 0.25μM - 5μM. (B) Inhibition of MukBF ATPase by MukE was dependent on the concentration of MukF. Concentrations of MukB and MukE were constant with MukB, 0.5μM, MukE 2.5μM while concentration of MukF ranged between 0.25μM - 5μM.

**Figure 7 – figure supplement 1.**
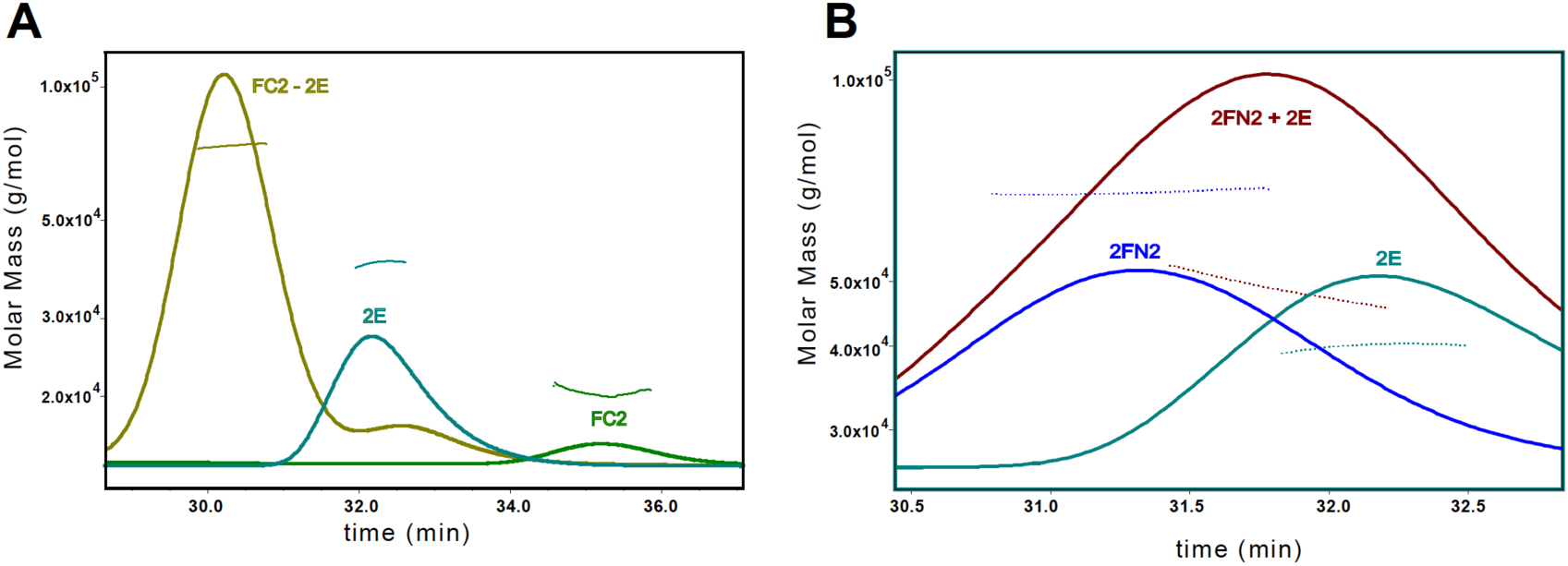
SEC-MALS analysis of MukE binding to FC2 and FN2. The samples contained a mixture of (A) FC2 and MukE at molar ratio of 1:1, monomer to dimer, **B**: FN2 and MukE at ratio 1:1.5 dimer:dimer.

**Figure 8 – figure supplement 1.**
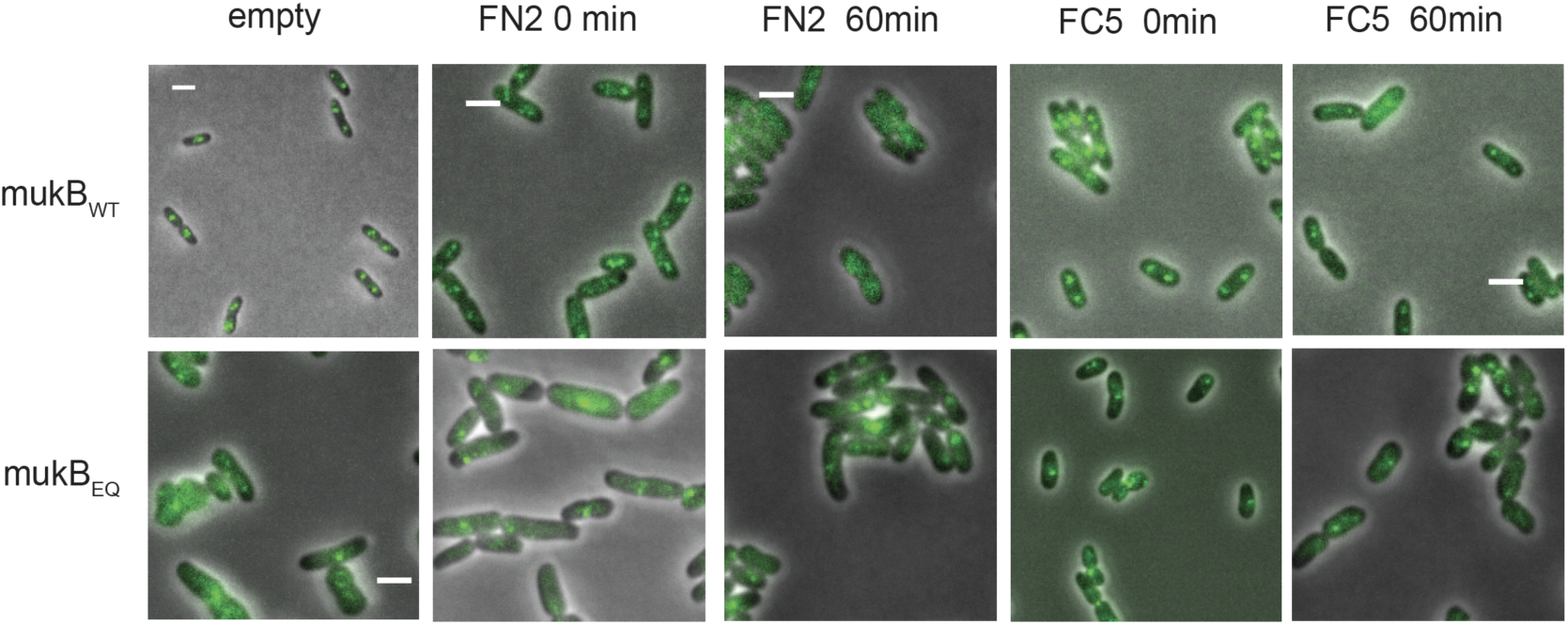
MukBEF foci in cells carrying either *mukB* wt or *mukB*^*EQ*^ mYPet chromosomal genes, before and after 1hr overexpression of MukF N- and C-terminal domain fragments, FN2 and FC5.

## References

ATP hydrolysis is required for cohesin’s association with chromosomes. P Arumugam, S Gruber, K Tanaka, CH Haering, K Mechtler, K Nasmyth (2003) Current Biology 13:1941–53. http://doi.org/10.1016/j.cub.2003.10.036

Cohesin’s ATPase activity is stimulated by the c-terminal winged-helix domain of its kleisin subunit P Arumugam, T Nishino, CH Haering, S Gruber, K Nasmyth (2006) Current Biology 16:1998–2008, http://dx.doi.org/10.1016/j.cub.2006.09.002

In vivo architecture and action of bacterial structural maintenance of chromosome proteins A Badrinarayanan, R Reyes-Lamothe, S Uphoff, MC Leake,DJ Sherratt (2012a) Science 338:528–53 http://doi:10.1126/science.1227126

The Escherichia coli SMC complex, MukBEF, shapes nucleoid organization independently of DNA replication. A Badrinarayanan, C Lesterlin, R Reyes-Lamothe, D Sherratt (2012b) Journal of Bacteriology 194:4669–76. http://doi:10.1128/JB.00957-12.

Releasing activity disengages cohesin’s Smc3/Scc1 interface in a process blocked by acetylation. F Beckouët, M Srinivasan, MB Roig, KL Chan, JC Scheinost, P Batty, B Hu, N Petela, T Gligoris, AC Smith, L Strmecki, BD Rowland, K Nasmyth (2016) Molecular Cell 61:563–74. http://doi:10.1016/j.molcel.2016.01.026.

DomainSieve: a protein domain-based screen that led to the identification of Dam-associated genes with potential link to DNA maintenance P Brezellec, M Hoebeke, M-S Hiet, S Pasek J-L Ferat (2006) Bioinformatics 22:1935–1941. http://doi:10.1093/bioinformatics/btl336

Prophase pathway-dependent removal of cohesin from human chromosomes requires opening of the Smc3–Scc1 gate J Buheitel, O. Stemmann (2013) The EMBO Journal 32:666–676. http://doi:10.1038/emboj.2013.7

MukB-mediated catenation of DNA is ATP and MukEF independent S Bahng, R Hayama, KJ Marians (2016) Journal of Biological Chemistry 291:23999–24008, http://doi:10.1074/jbc.M116.749994

Eco1-dependent cohesin acetylation during establishment of sister chromatid cohesion. TR Ben-Shahar, S Heeger, C Lehane, P East, H Flynn, M Skehel, F Uhlmann (2008) Science 321:563–6. http://doi:10.1126/science.1157774.

An asymmetric SMC-kleisin bridge in prokaryotic condensin. F Bürmann, HC Shin, J Basquin, YM Soh, V Giménez-Oya, YG Kim, BH Oh, S Gruber (2013) Nature Structural and Molecular Biology 20:371–9. http://doi:10.1038/nsmb.2488

The ATPases of cohesion interface with regulators to modulate cohesin-mediated DNA tethering. G Çamdere, V Guacci, J Stricklin, D Koshland (2015) eLife 4:e11315. http://dx.doi.org/10.7554/eLife.11315

Cohesin’s DNA exit gate is distinct from its entrance gate and is regulated by acetylation K-L Chan, M B Roig, B Hu, F Beckouët, J Metson, K Nasmyth (2012)Cell 150:961–974. http://dx.doi.org/10.1016/j.cell.2012.07.028

MukB acts as a molecular clamp in DNA condensation Y Cui, ZM Petrushenko, VV Rybenkov (2008) Nature Structural and Molecular Bioclogy 15:411–418 http://doi:10.1038/nsmb.1410

Condensin structures chromosomal DNA through topological links. S Cuylen, J Metz, CH Haering (2011) Nature Structural and Molecular Biology 18:894–901. http://doi:10.1038/nsmb.2087.

MukB colocalizes with the oriC region and is required for organization of the two Escherichia coli chromosome arms into separate cell halves. O Danilova, R Reyes-Lamothe, M Pinskaya, D Sherratt, C Possoz (2007) Molecular Microbiology 65:1485–92. http://doi:10.1111/j.1365-2958.2007.05881.x

Structure of full-Length SMC and rearrangements required for chromosome organization. ML Diebold-Durand, H Lee, LB Ruiz Avila, H Noh, HC Shin, H Im, FP Bock, F Bürmann, A Durand, A Basfeld, S Ham, J Basquin, BH Oh, S Gruber (2017) Molecular Cell 67:334–347.e5. http://doi:10.1016/j.molcel.2017.06.010.

Disengaging the Smc3/kleisin interface releases cohesin from Drosophila chromosomes during interphase and mitosis CS Eichinger, A Kurze, RA Oliveira, K Nasmyth (2013) The EMBO Journal 32:656–665. http://doi:10.1038/emboj.2012.346

Cohesin releases DNA through asymmetric ATPase-driven ring opening AMO Elbatsh, JHI Haarhuis, N Petela N, C Chapard, A Fish A, PH Celie, M Stadnik, D Ristic, C Wyman, RH Medema, K Nasmyth K, BD Rowland (2016) Molecular Cell 61:575–88. http://doi:10.1016/j.molcel.2016.01.025

The MukF subunit of Escherichia coli condensin: architecture and functional relationship to kleisins R Fennell-Fezzie, SD Gradia, D Akey, JM Berger (2005) The EMBO Journal 24:1921–30. http://doi:10.1038/sj.emboj.7600680

The deviant ATP-binding site of the multidrug efflux pump Pdr5 plays an active role in the transport cycle C Furman, J Mehla, N Ananthaswamy, N Arya, B Kulesh, I Kovach, SV Ambudkar, J Golin (2013) Journal of Biological Chemistry 288:30420–30431. http://dx.doi.org/10.1074/jbc.M113.494682

Structural Insights into ring formation of cohesion and related Smc complexes T Gligoris, J Löwe (2016) Trends in Cell Biology 1231:680–93. http://doi:10.1016/j.tcb.2016.04.002

Closing the cohesin ring: structure and function of its Smc3-kleisin interface TG Gligoris, JC Scheinost, F Bürmann, N Petela, K-L Chan, P Uluocak, F Beckouet, S Gruber, K Nasmyth, J Löwe (2014) Science 346:963–967. http://dx.doi.org/10.1126/science.1256917

The role of MukE in assembling a functional MukBEF complex M Gloyd, R Ghirlando, A Guarné (2011) Journal of Molecular Biology 412:578–590. http://doi:10.1016/j.jmb.2011.08.009

MukE and MukF form two distinct high affinity complexes. M Gloyd, R Ghirlando, LA Matthews, A Guarné (2007) Journal of Biological Chemistry 282:14373–8. http://doi:10.1074/jbc.M701402200

Mechanistic determinants of the directionality and energetics of active export by a heterodimeric ABC transporter N Grossmann, AS Vakkasoglu, S Hulpke, R Abele, R Gaudet, R Tampé (2014) Nature Communications 5:5419. http://dx.doi.org/10.1038/ncomms6419

Evidence that loading of cohesin onto chromosomes involves opening of its SMC hinge S Gruber, P Arumugam, Y Katou, D Kuglitsch, W Helmhart, K. Shirahige, K Nasmyth (2006) Cell, 127, 523–537 https://doi.org/10.1016/j.cell.2006.08.048

Chromosomal cohesin forms a ring S Gruber, CH Haering, K Nasmyth (2003) Cell 112:765–777. http://doi.org/10.1016/S0092-8674(03)00162-4

The cohesin ring concatenates sister DNA molecules.CH Haering, AM Farcas, P Arumugam, J Metson, K Nasmyth (2008) Nature 454:297–301. http://doi:10.1038/nature07098.

Molecular architecture of SMC proteins and the yeast cohesion complex. CH Haering, J Löwe, A Hochwagen, K Nasmyth (2002) Molecular Cell 9:773–88.http://dx.doi.org/10.1016/S1097-2765(02)00515-4

Structure and stability of cohesin’s Smc1-kleisin interaction.CH Haering, D Schoffnegger, T Nishino, W Helmhart, K Nasmyth, J Löwe (2004) Molecular Cell 15:951–64. http://doi:10.1016/j.molcel.2004.08.030

The MukB-ParC interaction affects the intramolecular, not intermolecular, activities of topoisomerase IV. R Hayama, S Bahng, ME Karasu, KJ Marians (2013) Journal of Biological Chemistry 288:7653–61 http://doi:10.1074/jbc.M112.418087.

Physical and functional interaction between the condensin MukB and the decatenase topoisomerase IV in Escherichia coli. R Hayama, KJ Marians (2010) Proceedings of the National Academy of Sciences of the United States of America 107:18826–31. http://doi:10.1073/pnas.1008140107

Mutants defective in chromosome partitioning in E. coli. Hiraga S, Niki H, Imamura R, Ogura T, Yamanaka K, Feng J, Ezaki B, Jaffé A. (1991) Research in Microbiology 142:189–94. http://dx.doi.org/10.1016/0923-2508(91)90029-A

At the heart of the chromosome: SMC proteins in action T Hirano (2006) Nature Reviews Molecular Cell Biology 7:311–322. http://dx.doi.org/10.1038/nrm1909

Structural basis for allosteric cross-talk between the asymmetric nucleotide binding sites of a heterodimeric ABC exporter M Hohl, LM Hurlimann, S Bohm, J Schoppe, MG Grutter, E Bordignon, MA Seeger (2014) Proceedings of the National Academy of Sciences of the United States of America 111:11025–11030. http://dx.doi.org/10.1073/pnas.1400485111

ABC-ATPases, adaptable energy generators fuelling transmembrane movement of a variety of molecules in organisms from bacteria to humans IB Holland, M A. Blight (1999) Journal of Molecular Biology 293:381–399. http://dx.doi.org/10.1006/jmbi.1999.2993

ATP hydrolysis is required for relocating cohesin from sites occupied by its Scc2/4 loading complex. B Hu, T Itoh, A Mishra, Y Katoh, KL Chan, W Upcher, C Godlee, MB Roig, K Shirahige, K Nasmyth (2011) Current Biology 2:12–24. http://doi:10.1016/j.cub.2010.12.004.

Characterisation of a DNA exit gate in the human cohesin ring PJ Huis in ‘t Veld, F Herzog, R Ladurner, IF Davidson, S Piric, E Kreidl, V Bhaskara, R Aebersold, JM Peters (2014) Science 346:968–72, http://doi:10.1126/science.1256904

Molecular Basis of SMC ATPase activation: role of integral structural changes of the regulatory subcomplex ScpAB K Kamada, M Miyata, T Hirano (2013) Structure 21:581–594. http://dx.doi.org/10.1016/j.str.2013.02.016

The Smc5/6 Complex Is an ATP-Dependent Intermolecular DNA Linker. T Kanno, DG Berta, C Sjögren (2015) Cell Reports 12:1471–82. http://doi:10.1016/j.celrep.2015.07.048.

Wapl controls the dynamic association of cohesin with chromatin. S Kueng, B Hegemann, BH Peters, JJ Lipp, A Schleiffer, K Mechtler, JM Peters (2006). Cell 127:955–67. Epub 2006 Nov 16. http://doi:10.1016/j.cell.2006.09.0402008

Structural biochemistry of ATP-driven dimerization and DNA-stimulated activation of SMC ATPases. A Lammens, A Schele, KP Hopfner (2004) Current Biology 14:1778–82. http://doi:10.1016/j.cub.2004.09.044

Escherichia coli condensin MukB stimulates topoisomerase IV activity by a direct physical interaction. Y Li, NK Stewart, AJ Berger, S Vos, AJ Schoeffler, JM Berger, BT Chait, MG Oakley (2010) Proceedings of the National Academy of Sciences of the United States of America 107:18832–7. http://doi:10.1073/pnas.1008678107.

Identification of interacting regions within the coiled coil of the Escherichia coli structural maintenance of chromosomes protein MukB. Y Li, CS Weitzel, RJ Arnold, MG Oakley (2009) Journal of Molecular Biology 391:57–73. http://doi:10.1016/j.jmb.2009.05.070

Mechanistic diversity in ATP-binding cassette (ABC) transporters KP Locher (2016) Nature Structural and Molecular Biology 23:487–493 http://doi.10.1038/nsmb.3216

Biochemical reconstitution of topological DNA binding by the cohesin ring. Y Murayama, F Uhlmann (2014) Nature 505:367–71 http://doi:10.1038/nature12867.

Chromosome segregation: how to open cohesin without cutting the ring? Y Murayama, F. Uhlmann (2013) The EMBO Journal, 32, 614–6 http://doi:10.1038/emboj.2013.22.

DNA entry into and exit out of the Cohesin Ring by an interlocking gate mechanism Y Murayama, F Uhlmann (2015) Cell 163:1628–1640. http://dx.doi.org/10.1016/j.cell2015.11.030

Cohesin: a catenase with separate entry and exit gates? K Nasmyth (2011) Nature Cell Biology 13:1170–7. http://doi:10.1038/ncb2349

The new gene mukB codes for a 177 kd protein with coiled-coil domains involved in chromosome partitioning of E. coli. H Niki, A Jaffé, R Imamura, T Ogura, S Hiraga (1991) EMBO Journal 10:183–93.

The bacterial chromosome: architecture and action of bacterial SMC and SMC-like complexes. S Nolivos, D Sherratt (2013) FEMS Microbiology Reviews 38:380–92 http://doi:10.1111/1574-6976.12045

MatP regulates the coordinated action of topoisomerase IV and MukBEF in chromosome segregation. S Nolivos, AL Upton, A Badrinarayanan, J Müller, K Zawadzka, J Wiktor, A Gill, LK Arciszewska, E Nicolas, DJ Sherratt (2016) Nature Communications 28:10466 http://doi:10.1038/ncomms10466.

Kite Proteins: a Superfamily of SMC/Kleisin Partners Conserved Across Bacteria, Archaea, and Eukaryotes. JJ Palecek, S Gruber (2015) Structure 23:2183–90. http://doi:10.1016/j.str.2015.10.004.

Antagonistic interactions of kleisins and DNA with bacterial Condensin MukB ZM Petrushenko, C-H Lai, VV Rybenkov (2006) Journal of Biological Chemistry 281:34208–34217, http://doi:10.1074/jbc.M606723200

The mechanism of ABC transporters: general lessons from structural and functional studies of an antigenic peptide transporter. E Procko, ML O’Mara, WF Drew Bennett, DP Tieleman, R Gaudet (2017) The FASEB Journal 23:1287–1302 http://doi:10.1096/fj.08-121855

Mutational analysis of MukE reveals its role in focal subcellular localization of MukBEF. W She, E Mordukhova, H Zhao, ZM Petrushenko, VV Rybenkov (2013) Molecular Microbiology 87:539–552 http://doi:10.1111/mmi.12112

High-throughput, subpixel precission analysis of bacterial morphogenesis and intracellular spatio-temporal dynamics O Sliusarenko, J Heinritz, T Emonet, C Jacobs-Wagner (2011) Molecular Microbiology 80:612–27. http://doi:101111/j.1365-2958.2011.07579.x

Molecular basis for SMC rod formation and its dissolution upon DNA binding Y-M Soh, F Bürmann, H-C Shin, T Oda, KS Jin, CP Toseland, C Kim, H Lee, SJ Kim, M-S Kong, M-L Durand-Diebold, Y-G Kim, HM Kim, NK Lee, M Sato, B-H Oh, S Gruber (2015) Molecular Cell 57:290–303. http://dx.doi.org/10.1016/j.molcel.2014.11.023

Structural diversity of ABC transporters J ter Beek, A Guskov, DJ Slotboom (2014) The Journal of General Physiology 143:419–435. http://doi10.1085/jgp.201411164

The condensing complex is a mechanochemical motor that translocates along DNA T Terkawa, S Bisht, J Eeftens, C Deeker, C Haering, E Greene (2017) bioRxiv http://dx.doi.org/10.1101/137711

SMC complexes: from DNA to chromosomes F Uhlmann (2016) Nature Reviews Molecular Cell Biology 17:399–412, http://doi:10.1038/nrm.2016.30.

A molecular determinant for the establishment of sister chromatid cohesion. E Unal, JM Heidinger-Pauli, W Kim, V Guacci, I Onn, SP Gygi, DE Koshland (2008) Science 321:566–9. http://doi:10.1126/science.1157880.

Structural basis for the MukB-topoisomerase IV interaction and its functional implications in vivo.SM Vos, NK Stewart, MG Oakley, JM Berger (2013) EMBO Journal 32:2950–62. http://doi:10.1038/emboj.2013.218.

A repeated coiled-coil interruption in the Escherichia coli condensin MukB. CS Weitzel, VM Waldman, TA Graham, MG Oakley (2011) Journal of Molecular Biology 414:578–95. https://doi:10.1016/j.jmb.2011.10.028

Evolution of condensin and cohesin complexes driven by replacement of Kite by Hawk proteins. JN Wells, TG Gligoris, KA Nasmyth, JA Marsh (2017) Current Biololgy 27: R17–R18. http://doi:10.1016/j.cub.2016.11.050.

SMC condensin entraps chromosomal DNA by an ATP hydrolysis dependent loading mechanism in Bacillus subtilis. L Wilhelm, F Bürmann, A Minnen, HC Shin, CP Toseland, BH Oh, S Gruber (2015) elife 4:e06659. http://doi:10.7554/eLife.06659.

Structural studies of a bacterial condensin complex reveal ATP-dependent disruption of intersubunit interactions. Woo JS, Lim JH, Shin HC, Suh MK, Ku B, Lee KH, Joo K, Robinson H, Lee J, Park SY, Ha NC, Oh BH (2009) Cell 136:85–96. http://doi:10.1016/j.cell.20089.10.050

Complex formation of MukB, MukE and MukF proteins involved in chromosome partitioning in E. coli M Yamazoe, T Onogi, Y Sunako, H Niki, K Yamanaka, T Ichimura, S Hiraga (1999) EMBO Journal 18, 5873–5884, http://doi:10.1093/emboj/18.21.5873

Identification of two new genes, mukE and mukF, involved in chromosome partitioning in Escherichia coli. K Yamanaka, T Ogura, H Niki, S Hiraga (1996) Molecular and General Genetics 250:241–5.

ATP binding to the first nucleotide binding domain of multidrug resistance-associated protein plays a regulatory role at low nucleotide concentration, whereas ATP hydrolysis at the second plays a dominant role in ATP-dependent leukotriene C4 transport. R Yang, L Cui, Y.-x. Hou, JR Riordan, X.-b. Chang (2003) Journal of Biological Chemistry 278:30764–30771. http://dx.doi.org/10.1074/jbc.M304118200

Toward Determining ATPase Mechanism in ABC Transporters: Development of the Reaction Path-Force Matching QM/MM Method Y Zhou, P Ojeda-May, M Nagaraju, J Pu (2016) Methods of Enzymology 577:185–212. http://doi:10.1016/bs.mie.2016.05.054

